# Geographic isolation enables recurrent pollinator shifts despite hybridisation in the Cape’s hyperdiverse heathers

**DOI:** 10.1101/2023.07.19.549682

**Authors:** Seth D. Musker, Michael D. Pirie, Nicolai M. Nürk

## Abstract

Deciphering the ecological and geographic factors that influence the dynamics of population divergence can aid in understanding why some groups of organisms diversify more prolifically than others. One such diverse group is the heathers (*Erica*, Ericaceae), whose exceptional species richness in the Cape Floristic Region is enigmatic. Here, we study *Erica abietina*, a small but highly variable species complex with four described subspecies differing in geographic range, habitat, and floral characters associated with pollination. To understand the factors and forces that shaped its evolution, we evaluate the status of the subspecies and test for hybridisation, introgression, pollinator-driven divergence, and geographic population structure using genotyping-by-sequencing on samples across the entire distribution. We find that the four subspecies form variably distinct genetic groups, however, the most widespread subspecies exhibits cryptic diversity comprising two independent lineages that are geographically isolated and occur on different soil types. Phylogenetic results suggest that shifts between bird- and insect-pollination syndromes have occurred twice independently, with accompanying genetic divergence. However, for one pair of genetically distinct populations (*F*_*ST*_ *≈* 0.06) with different pollinators we uncover several individuals of hybrid origin at a site where they occur sympatrically. Together, these results suggest that floral differentiation driven by divergent selection acts in concert with geographic isolation to maintain reproductive isolation. Finally, we show that a reticulate history involving “ghost” introgression best explains the group’s evolution. Our results reveal a highly dynamic system whose diversity has been shaped by a variety of interacting forces, and we suggest that such systems are likely to have contributed substantially to the diversity of *Erica* and the Cape flora in general.

## Introduction

Patterns of biological diversity are shaped by a variety of geographic, ecological, and genetic factors. At the interface of micro- and macro-evolution, these factors interact in multifaceted ways to promote or inhibit population divergence and eventual speciation (Templeton, 1981). This means that for any group of organisms, teasing apart the relative importance of the factors driving diversification requires combining genetic data with information on the group’s distribution, ecology, and environment (Via, 2009; Donoghue and Sanderson, 2015). This can be challenging in highly dynamic systems—those in which species limits are unclear and a variety of processes, such as hybridisation, introgression, genetic drift, and selection, may act together. Such dynamic systems should be prevalent in global biodiversity hotspots (Myers et al., 2000), which are often typified by high rates of speciation (e.g., Madriñán et al., 2013). The Cape Floristic Region (CFR) is one such hotspot, hosting a megadiverse flora of over 9000 vascular plant species of which a remarkable 70% are endemic (Manning and Goldblatt, 2012). Considerable research effort has focused on the macroevolutionary factors responsible for the CFR’s exceptional diversity (Verboom et al., 2009). In contrast, relatively few studies have investigated the dynamics of species divergence in its early stages using genomic methods and representative population sampling (cf. Lexer et al., 2014; Prunier and Holsinger, 2010; Prunier et al., 2017). Here, we present the first genomic study of highly dynamic early-stage diversification in *Erica* L. (Ericaceae), which, with almost 700 species, is easily the largest genus in the CFR (Manning and Goldblatt, 2012).

The presence of *Erica* species is a defining feature of the fynbos vegetation that dominates much of the CFR (Bergh et al., 2014). *Erica* species also contribute substantially to several distinctive features of the fynbos, including a proclivity for extremely nutrient-poor soils (Bradshaw and Cowling, 2014) and exceptionally high species turnover at small spatial scales (Kruger and Taylor, 1980). The genus owes its richness to rapid and prolific diversification that evidently began upon its arrival in the CFR around 10-15 million years ago (Ma) (Pirie et al., 2016, 2019). However there remains considerable uncertaintly regarding the factors that drove the Cape clade radiation. One suggestion is that frequent pollinator shifts, which have contributed to plant diversification in the CFR in general (e.g., Johnson, 1996; van der Niet and Johnson, 2009), may have done the same in *Erica*. This is supported by evidence that floral traits are exceptionally labile throughout the phylogeny of the Cape *Erica* clade (Pirie et al., 2011). In particular, shifts between insect- and bird-pollination may have been especially influential (van Der Niet et al., 2014). At least sixty *Erica* species are thought to be pollinated by birds, the vast majority of which are confined to the CFR (Rebelo et al., 1985), and almost all are pollinated by the CFR-endemic Orange-breasted Sunbird (*Anthobaphes violacea*; Rebelo et al., 1984; Coetzee, 2016; Coetzee et al., 2020). Sunbird-pollinated *Erica* tend to have long (18-26 mm) corolla tubes that match the dimensions of the birds’ bills (Rebelo et al., 1984, 1985; Barnes et al., 1995), whereas most insect-pollinated species have short corollas (Rebelo et al., 1984). Bird-pollinated *Erica* are also unscented and come in a variety of flower colours, whereas insect-pollinated species are often sweetly scented and typically have pink or white flowers (Rebelo et al., 1984, 1985; Oliver and Oliver, 2002, 2005). Notably, despite the distinctness of these pollination syndromes (Rebelo et al., 1985) they seem to be very weakly conserved among closely related species (Pirie et al., 2011). What remains unclear, however, is how and to what degree such pollinator shifts might have stimulated speciation and contributed to Cape *Erica* diversity, and whether additional factors, such as geographic isolation and other forms of niche divergence, played supporting or even superior roles.

Here, we focus on the *Erica abietina* species complex, which comprises several phenotypically distinct forms varying in corolla tube length and flower colour (Table 1). On the basis of this floral variation the species has been classified into four subspecies (Oliver and Oliver, 2002; Pirie et al., 2017), all of which are confined to the Cape Peninsula (CP; Fig. 1). The CP is a 50 × 15 km mountain range of rugged topography at the south-western tip of South Africa that extends from Table Mountain in the north to Cape Point in the south. Being largely surrounded by ocean, the CP is isolated from the rest of the Cape Fold Mountains by an extensive sandy plain (> 40 km wide) whose low elevation placed it below sea level during Pleistocene interglacials (Adamson, 1959), and which now holds the city of Cape Town. The CP is a hotspot within a hotspot of floristic diversity (Cowling et al., 1996) with over 2200 plant species (Trinder-Smith et al., 1996) of which 158 (including *≥* 39 *Erica* species) are endemic (Helme and Trinder-Smith, 2006). All of this implies that *E. abietina* may be experiencing diversifying forces in a system akin to a “continental archipelago” (cf. Hughes and Eastwood, 2006), making it an ideal system for studying the dynamics of the early stages of species diversification.

**Table 1.** Characteristics of the four subspecies of *Erica abietina*.

| Subspecies | Corolla length | Corolla colour | Floral scent | Sepals | Ovary | Range |
| --- | --- | --- | --- | --- | --- | --- |
| *E. a. abietina* | 18–26 mm | Crimson to dark red | None | Lanceolate-attenuate/ acuminate; pilose | Elongate obovoid; pubescent | Northern CP: TM high plateau, N and W slopes |
| *E. a. atrorosea* | 18–22 mm | Rose to deep rose pink | None | Lanceolate-acute; glabrous to sparsely puberulous | Ellipsoid; glabrous | Widespread on CP: E slopes of TM S to Cape Point |
| *E. a. constantiana* | 8–11 mm | Pale to deep rose pink | Sweet, lemony | Lanceolate-attenuate/ acuminate; glabrous to sparsely puberulous | Squat obovoid; puberulent | Central CP: S slopes of TM to Chapman's Peak |
| *E. a. diabolis* | 11–14 mm | Rose pink | None | Lanceolate-attenuate/ acuminate; pilose | Obovoid; pubescent | Northern CP: Devil's Peak |

CP: Cape Peninsula; TM: Table Mountain.

**Figure 1.**
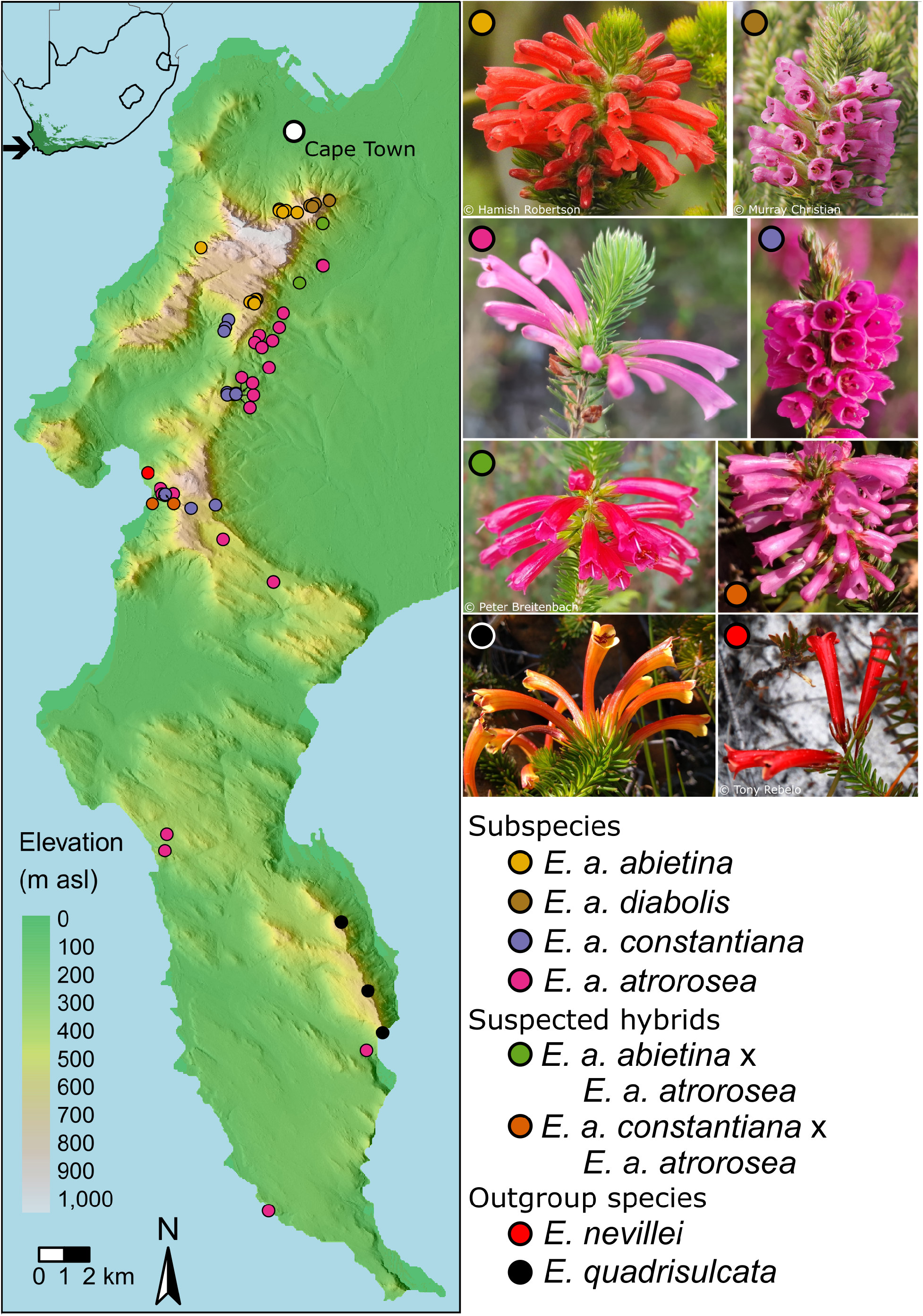
*Left*: Map of the Cape Peninsula showing elevation and hillshade with localities of collected samples. *Inset*: Map of South Africa showing the extent of the Fynbos biome and location of the Cape Peninsula. *Right*: Images of the focal taxa including putative hybrids.

To illuminate the evolutionary history of the *E. abietina* species complex, we use genotyping-by-sequencing (GBS; Elshire et al., 2011) coupled with population-genetic and phylogenetic analyses. Firstly, we examine whether genetic differentiation in the species complex is correlated with geography and to what extent it corroborates current taxon boundaries. Secondly, we infer phylogenetic relationships between individuals and populations to test whether shifts between bird and insect pollination syndromes have occurred multiple times, so contributing to lineage divergence. Finally, we test for evidence of ongoing and ancient gene flow in order to infer the influence of hybridisation and introgression on species divergence, both past and present.

## Materials and Methods

### Sample collection and sequencing

We collected fresh leaf material from at least six individuals of each formally recognised taxon in addition to several individuals of putative hybrid origin (Table 1) and attempted to sample from across the full geographic range of each taxon (Fig. 1). We followed the taxon concepts (Meier, 2017) outlined by Oliver and Oliver (2002) and refined by Pirie et al. (2017) when identifying specimens. In addition, we sampled from the two putatively closest outgroups of *E. abietina, E. nevillei* (one sample) and *E. quadrisulcata* (three samples), both of which are endemic to the Cape Peninsula (Pirie et al., 2017). DNA extraction in *Erica* is challenging (Bellstedt et al., 2010), and we adapted the protocol of Inglis et al. (2018) to retrieve sufficient DNA for sequencing (see Supplemetary Material).

GBS library preparation and sequencing was done by Novogene Genome Sequencing Company Ltd. (Beijing, China).

Protocols followed the original Elshire et al. (2011) pipeline except that two restriction enzymes were used: MseI and HaeIII. Libraries were paired-end sequenced in two separate batches using an Illumina NovaSeq 6000 instrument to produce 144 bp-long reads after barcode removal. The two batches differed in the amount of sequencing effort per sample, though we found this to have little impact on the quality of the results. One sample was included in both batches to test the effect of sequencing effort on genotype calling accuracy, which we found to be negligible. Full details are available in the Supplemetary Material.

Raw reads were quality-checked with FastQC v.0.11.9 (http://www.bioinformatics.bbsrc.ac.uk/projects/fastqc) and MultiQC v.1.14 (Ewels et al., 2016). Adapter removal, quality trimming and filtering of reads was done with fastp v.0.20.1 (Chen et al., 2018, parameters: *–overrepresentation_analysis –trim_poly_g –qualified_quality_phred 20 –unqualified_percent_limit 30 –average_qual 20 –length_required 100*). Clean reads were aligned to a draft reference genome of *Erica cerinthoides* (Musker et al., unpublished) using BWA-MEM v.0.7.17-r1198-dirty (Li, 2013) with default parameters, followed by alignment sorting and indexing using SAMtools v.1.11-2 (Danecek et al., 2021) and merging using BamTools v.2.1.1 (Barnett et al., 2011).

Variants were called for all samples simultaneously using Freebayes v.1.3.4 (Garrison and Marth, 2012, parameters: *–min-base-quality 3 –min-mapping-quality 10 –skip-coverage 10000 –use-best-n-alleles 4*). Table S1 outlines the variant calling results and the effects of quality filtering. Genotypes with read depth < 3 were recoded as missing using vcftools v.0.1.17 (Danecek et al., 2011). Using BCFTools v.1.12-5 (Danecek et al., 2021) we then kept variants with missingness (proportion of missing genotypes) < 0.5, minor allele frequency (MAF) > 0.01, allele balance < 0.01 or between 0.25 and 0.75, and MQM/MQMR (ratio of forward to reverse read mapping qualities) between 0.9 and 1.1. We then used *vcfallelicprimitives* from vcflib v.1.0.1 (Garrison, 2012) to decompose complex variants into SNPs where possible, and then removed non-SNP variants and non-biallelic SNPs. We used the BCFTools *fill-tags* plugin to test for excess heterozygosity, and removed SNPs with a p-value *≤* 0.2. We applied further filters as appropriate and used a naming scheme based on these filters (see Table S1). Filtering based on linkage disequilibrium (LD) was done using PLINK v.1.90b6.21 (Purcell et al., 2007, parameters: *–indep-pairwise 50 5 0*.*2*).

For calculating sequence diversity and absolute sequence divergence as well as for individual-level phylogeny inference, we generated an “all sites” VCF file by supplementing the SNP_m10 set (SNPs-only, missingness < 10%; Table S1) with invariant sites. This was done by re-running FreeBayes as above but adding the “*–report-monomorphic*” switch, followed by removing sites with missingness > 0.1, decomposing complex variants as above, keeping only invariant sites, and finally combining the invariant sites with the SNP_m10 set using BCFtools. Several of the above processes were parallelised using GNU Parallel v.20200522 (Tange, 2020). The majority of the following analyses were conducted in the R statistical environment (R Core Team, 2021) using R v.4.1.2. VCF files were read using vcfR v.1.13.0 (Knaus and Grünwald, 2017).

### Analysis of population structure

To assess population structure without any prior assumptions about group membership, we employed three complementary methods. Firstly, we ran a Principal Component Analysis on the SNP_m10_LD_maf04 set using the adegenet v.2.1.5 (Jombart and Ahmed, 2011) function *glPca*. Secondly, we employed the admixture model (Pritchard et al., 2000) for values of *K* (the number of ancestral populations) ranging from 2 to 8 using the sparse nonnegative matrix factorization (sNMF) algorithm (Frichot et al., 2014) implemented in LEA v.3.6.0 (Frichot and François, 2015), with default parameter values. For this we used the SNP_m10_LD_maf04 set and excluded the outgroups *E. nevillei* and *E. quadrisulcata*. For each *K* we ran 100 independent repetitions of the algorithm and summarised the outputs using the CLUMPAK method (Kopelman et al., 2015) implemented in STARMIE v.0.1.3 (https://github.com/sa-lee/starmie). We generated bar plots of ancestry proportions using pophelper v.2.3.0 (Francis, 2017). Lastly, to better visualise the connections between individuals as well as explore the hierarchical structure of genetic variation, we employed network analysis using NetView v.1.0 (Steinig et al., 2016, https://github.com/esteinig/netview). With the ALL_m10 set, we used pixy v.1.2.5.beta1 (Korunes and Samuk, 2021) to calculate the harmonic mean of absolute sequence divergence (*dxy*) between individuals to generate the genetic distance matrix required by NetView. We ran the network inference algorithm with values of *k* (the maximum number of mutual nearest neighbours) of 5, 10, 15 and 20, each time also estimating the minimum spanning tree to ensure a connected network was returned. We visualised each network with iGraph v.1.3.5 (Csardi and Nepusz, 2006) using the Kamada-Kawai spring-based layout algorithm (Kamada et al., 1989) to make the lengths of the connecting edges proportional to their associated genetic distance, and coloured the edges based on whether they were unique to the minimum spanning tree.

Based on these analyses we identified two genetically and geographically distinct northern and southern clusters within *E. a. atrorosea* (see Results), which we refer to as *E. a. atrorosea* (North) and *E. a. atrorosea* (South). We distinguished these clusters for further analyses that required individuals to be grouped.

### Detecting recent hybrids

To investigate the presence of hybrids between genetically distinct clusters, we sequenced four individuals of putative hybrid origin which we identified based on a combination of morphological features and geographic location (Fig. 1). Firstly, we identified two individuals from the mid-elevation eastern slopes of Table Mountain (TM) from populations showing a range of intermediate flower colours (magenta to cerise) between *E. a. abietina* (light red; TM plateau) and *E. a. atrorosea* (North) (pink; TM lower eastern slopes). Secondly, we identified two individuals from Blackburn Ravine with floral tube lengths intermediate between *E. a. atrorosea* (South) (18–22 mm) and *E. a. constantiana* (8–11 mm). To test the hybrid origin of these individuals we employed NewHybrids v.2.0 (Anderson, 2008) in combination with sNMF. For each set of putative parents and hybrids we subset the SNP_m10_LD_maf04 set to only the relevant individuals, removed resulting monomorphic sites, and ran sNMF with *K* = 2, again repeating the algorithm 100 times and summarising ancestry coefficients across runs as above. We then identified putatively non-admixed individuals as those whose maximum individual ancestry coefficient was *≥* 0.9, setting these as “P0” or “P1” in NewHybrids. To maximise the information content of the SNPs used, we calculated per-SNP *F*_*ST*_ (Weir and Cockerham, 1984) between P0 and P1 using the HierFstat v.0.5-11 (Goudet, 2005) *basic*.*stats* function and chose the 500 SNPs with the highest *F*_*ST*_ while also only keeping one SNP per contig to avoid linkage effects. We used DartR v.2.0.4 (Gruber et al., 2018) to convert the data into NewHybrids format. We then ran NewHybrids for 150,000 MCMC iterations, discarding the first 50,000 as burn-in. Based on these analyses we identified individuals showing evidence of recent hybrid origin (including backcrosses). To refer to data sets excluding these individuals, we append the suffix “_noHybrids”.

### Phylogenetic analysis

We inferred two individual-level phylogenies, one including all individuals (n = 65) and one excluding putative hybrid individuals (n = 45). We used IQ-TREE v.2.0.6 (Minh et al., 2020) using the full concatenated alignments. The ALL_m10 and ALL_m10_noHybrids VCF files were converted to fasta format using VCF2PHYLIP v.2.0 (Ortiz, 2019). For each analysis we chose the best-fitting substitution model using ModelFinder (Kalyaanamoorthy et al., 2017) and the default Bayesian Information Criterion (BIC). We estimated node certainty with 1000 ultrafast bootstrap replicates (Hoang et al., 2018) and 1000 SH approximate likelihood ratio test (SH-alrt; Guindon et al., 2010) replicates, and ran four independent runs to improve the search of the likelihood space. We plotted the maximum-likelihood tree with GGTREE v.3.2.1 (Yu et al., 2017) after rooting the tree at *E. quadrisulcata* with ape v.5.0 (Paradis and Schliep, 2019).

### Summary statistics

#### Individual-level summary statistics

To quantify genetic diversity we calculated per-sample multi-locus heterozygosity (MLH) using the R package INBREEDR v.0.3.3 (Stoffel et al., 2016) with the SNP_m10_LD_maf04 set. This revealed three anomalous samples within *E. abietina*, two with unusually low MLH which may be inbred (one from *E. a. abietina* and one from *E. a. atrorosea* [North]), and one with unusually high MLH (from *E. a. diabolis*) which may have been a chimeric sample stemming from a collection mishap in which two adjacent individuals were assumed to be one (Fig. S1).

#### Isolation-by-distance

To test for genetic isolation-by-distance (IBD) between individuals within taxa, Mantel tests (Mantel, 1967) were performed separately for each taxon and for the two clusters within *E. a. atrorosea*. Geographic distances were determined after projecting collection localities onto the Hartebeesthoek94 coordinate reference system, and *dxy* was used as the genetic distance. Mantel tests were performed using the R package VEGAN v.2.6-4 (Oksanen et al., 2019), testing for statistical significance using 1000 permutations.

#### Grouping individuals into populations

Confidently estimating population-level summary statistics typically necessitates grouping individuals into putatively monophyletic, panmictic and outbred “populations”. Based on the previous analyses, we removed 17 individuals with potentially mixed ancestry and three with anomalous MLH, and assigned the remainder to five distinct populations within *E. abietina*. Finally, we split *E. a. atrorosea* (South) into two groups, creating a separate group for three individuals collected from Blackburn Ravine whose inclusion would make the population paraphyletic according to the phylogenetic analyses (see Fig. 5). We distinguish these populations from the previously named entities as described in Table 2. All of the analyses described below used this assignment scheme.

**Table 2.** Assignment of taxa to newly defined populations for group-level analyses.

| Taxon | Population | *n* |
| --- | --- | --- |
| *E. a. abietina* | *ABI* | 11 |
| *E. a. atrorosea* (South) | *ATRO<sub>S</sub>* | 7 |
| *E. a. atrorosea* (North) | *ATRO<sub>N</sub>* | 7 |
| *E. a. atrorosea* (Blackburn Ravine) | *ATRO<sub>BLACKBURN</sub>* | 3 |
| *E. a. constantiana* | *CONS* | 8 |
| *E. a. diabolis* | *DIAB* | 5 |

#### Population-level summary statistics

To estimate the magnitude of pairwise genomic differentiation between the five populations plus the two outgroup taxa, we calculated pairwise Hudson’s *F*_*ST*_ (Hudson et al., 1992; Bhatia et al., 2013) using ADMIXTOOLS2 (Maier et al., 2022).

#### Investigating population history

We next aimed to investigate the branching history of the group and test hypotheses of ancient introgression between populations. To estimate the population phylogeny while accounting for incomplete lineage sorting, we used a polymorphism-aware model (PoMo; Schrempf et al. 2019) implemented in IQ-TREE v.2.0.6 (Minh et al., 2020) with the SNP_m10_maf01 set. We set the substitution model to GTR+G4 (the GTR model [Tavaré 1986] with four discrete gamma rate categories), conducted 1000 ultrafast bootstrap replicates, and left all other parameters at their default values.

To search for evidence of ancient introgression, we used the *findGraphs* method implemented in ADMIXTOOLS2 (Maier et al., 2022) which conducts a heuristic search of the graph space by iteratively proposing modifications to the current graph – such as modifying the tree topology and adding admixed edges – each time evaluating the new graph’s likelihood score, which is determined by comparing the f3-statistics (Peter, 2016) predicted by the graph to those estimated from the data. The algorithm attempts to find the best-fitting graph for the lowest number of allowed admixed edges (*N*_*ADMIX*_) before adding more admixed edges one at a time. We ran this analysis in two ways: first by specifying the population tree estimated by PoMo as the starting tree, and second by setting *E. quadrisulcata* as the outgroup without specifying any other restrictions to the randomly generated starting tree. In each case we ran the analysis five times. We started the graph searches at *N*_*ADMIX*_ = 0 and set the maximum value of *N*_*ADMIX*_ to 2. To search the likelihood space more exhaustively, we set the total number of generations after which to stop to 5000; the number of generations without improvement after which to stop to 100; the number of graphs evaluated in each generation to 30; and the *plusminus_generations* parameter (which helps to break out of local optima) to 20. All other parameters were kept at their default values. We manually inspected the ten resulting best graphs for each value of *N*_*ADMIX*_ to check for concordance in the graph topology, edge weights, and admixture proportions.

We compared alternative best-fitting graphs in a pairwise manner using a resampling procedure implemented in ADMIXTOOLS2 v.2.0.0 and described in Maier et al. (2022), which aims to test whether two graphs have similar predictive power. We used *qpgraph_resample_multi* to generate 100 replicate bootstrap resampled SNP block training and test sets and then evaluate each graph by estimating its weights using the training set and calculating its “out-of-sample” likelihood score using the (unseen) test set. This procedure allows to test the null hypothesis that the differences between two graphs’ out-of-sample scores for each bootstrap replicate are equal to zero, i.e., that both graphs have equivalent predictive power. This was done using *compare_fits*, which conducts a two-sided *z*-test on the score differences, assuming a normal distribution of values and known standard deviation.

## Results

### Read mapping

After read mapping, a mean of 67.4% (±2.94% SD) of read pairs mapped properly to the *E. cerinthoides* draft genome. There was a small but statistically significant difference in mapping rate between the two batches (batch 1: 65.1%, batch 2: 67.9%; linear model, F(1,64) = 9.84, p = 0.00258) suggesting there were more duplicated reads in batch 2. Mean mapping depth was 5.62 (±0.50 SD) and 9.10 (±1.63 SD) for batches 1 and 2, respectively.

### Variant calling

Variant calling followed by stringent quality filtering resulted in a rich and highly complete SNP data set (Table S1). Individuals sequenced in batch 1 had more missing genotypes in the SNP_m10 set (mean = 8.81%, SD = 3.09%) than batch 2 individuals (mean = 2.50%, SD = 1.86 %), while the highest missingness in any individual was 13.4%. Genotype calls had lower read counts for batch 1 individuals (mean = 10.3, range = 8.1–12.7) than batch 2 individuals (mean = 22.6, range = 12.5–32.5). Missingness was generally higher in the outgroups (*E. nevillei* : 9.14%; *E. quadrisulcata*: 7.01–11.2%), though this was not appreciable, suggesting that allele dropout did not affect library preparation or variant detection.

### Population structure

The Mantel tests for isolation-by-distance indicated that geographic distance was significantly correlated with genetic relatedness for all taxa, though the strength of IBD, measured as Mantel’s *r*, was variable (Table 3). All analyses of population structure indicated strong genetic clustering corresponding broadly to the current taxonomic treatment of the species complex, with some noteworthy exceptions. The PCA eigenvalues (Fig. 2, *inset*) exhibited a steep decline in explained variance from axes 1 to 4 followed by a plateau from axes 5 to 7, after which they declined gradually. This pattern suggested the existence of either five or eight clusters in the data, as *n −* 1 axes are required to distinguish *n* clusters. In contrast, and despite the exclusion of outgroups, the sNMF-based cross-entropy criterion suggested *K* = 1 or *K* = 2 to be optimal given a 5% genotype masking rate, although higher values of *K* generally recovered sensible groupings of individuals in concordance with the PCA and NetView analyses (Figs. 3,S2).

**Table 3.** Results of the Mantel tests of isolation-by-distance within taxa.

| Population | Mantel's $r$ | $p$-value | No. perm. |
| --- | --- | --- | --- |
| *E. a. abietina* | 0.686 | 0.001 | 999 |
| *E. a. atrorosea* (South) | 0.449 | 0.005 | 999 |
| *E. a. atrorosea* (North) | 0.876 | 0.001 | 999 |
| *E. a. constantiana* | 0.718 | 0.001 | 999 |
| *E. a. diabolis* | 0.573 | 0.021 | 719 |

**Figure 2.**
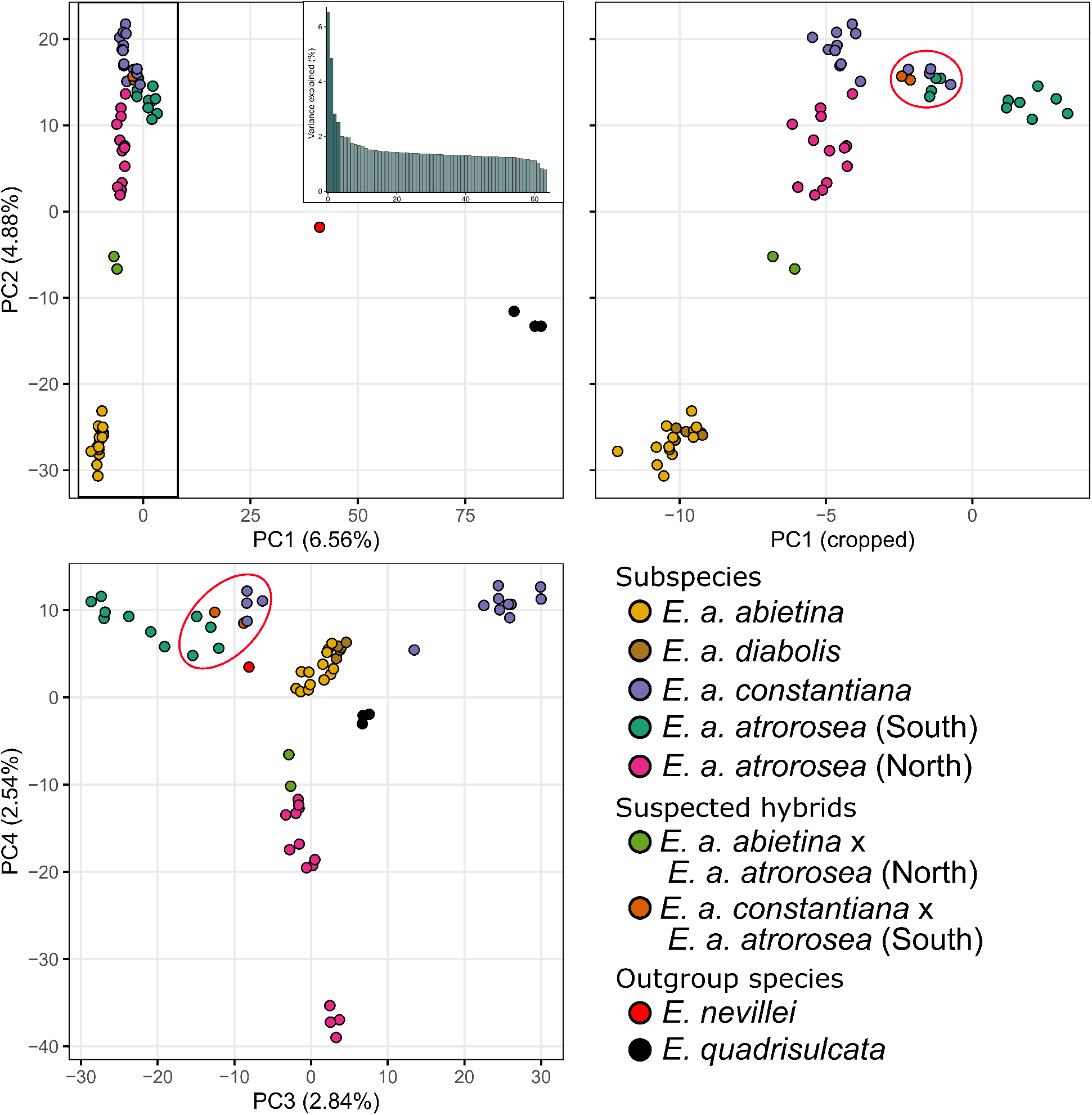
Ordination of the genotype matrix by PCA. The box in the first plot indicates the area covered by the second plot. Samples from Blackburn Ravine are indicated by red ellipses. The inset shows a plot of the variance explained by each PCA axis (also shown in brackets in axis titles). Note especially the close relationship between *E. a. abietina* and *E. a. diabolis* and their distinctness from the other subspecies; the positioning of the two samples identified as *E. a. abietina x E. a. atrorosea* hybrids based on morphology; and the positioning of samples from Blackburn Ravine.

**Figure 3.**
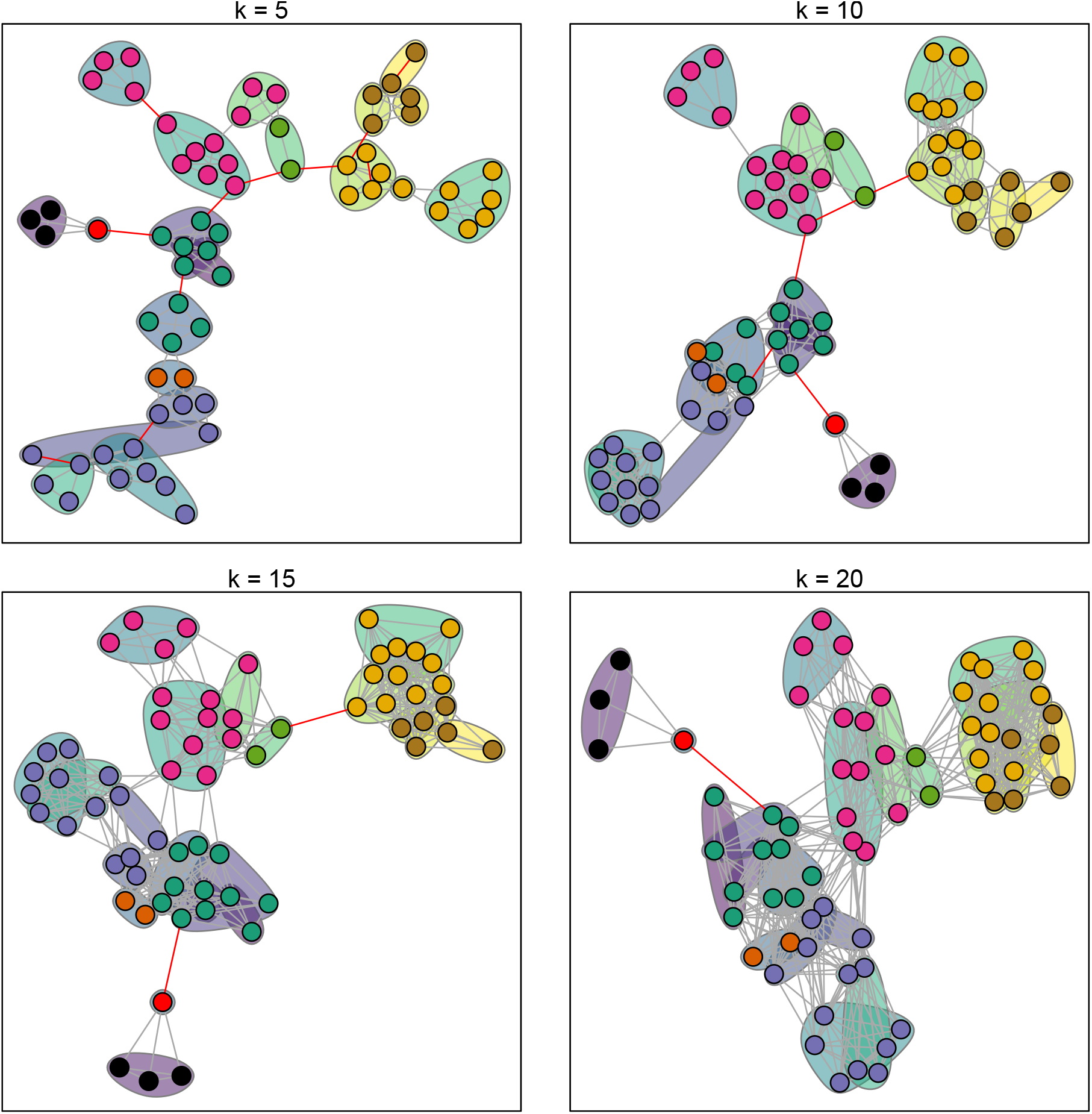
Genetic distance network visualisation. Panels show the NetView networks for different values of *k* (no. of allowed connections per node) based on *dxy*. Nodes represent individuals and colours reflect prior population assignments (Fig. 2). The shaded regions envelope individuals located geographically close to each other, with the shading representing mean latitude (Viridis colour scale; lighter colours are more northerly). Red edges are unique to the minimum spanning tree. Note how the finer scale differences are more noticeable for small values of *k*, becoming obscured by the broader patterns as *k* increases.

The first principal component axis (PC1; Fig. 2) primarily distinguished the outgroups from *E. abietina* and showed *E. nevillei* to be the closer of the two outgroups. Within *E. abietina*, PC2 and, to a lesser extent, PC1 distinguished *E. a. abietina* plus *E. a. diabolis* from the rest of the subspecies, and the sNMF results also recovered these two groups as the most important ancestral clusters at *K* = 2 (Fig. S2). Notably, there was consistent support for two distinct genetic clusters within *E. a. atrorosea*: One group (*E. a. atrorosea* [South]) consisted of individuals from the southern parts of the Cape Peninsula ranging from Cape Point to Silvermine, while the other (*E. a. atrorosea* [North]) comprised northern individuals collected along the lower eastern slopes of Table Mountain. In PCA space, *E. a. atrorosea* (North) fell between the two major clusters while still being closer to *E. a. atrorosea* (South) and *E. a. constantiana*. The NetView analysis (Fig. 3) revealed more fine-scale patterns of genetic structure and relatedness, especially at lower values of *k* (the maximum number of connections allowed between individuals). Within *E. a. abietina*, individuals sampled from different parts of Table Mountain were clearly recovered as belonging to distinct network clusters at *k* = 5. Within *E. a. atrorosea* (North), the four southernmost individuals were consistently recovered as distinct from and unconnected to any other populations, unlike the rest of *E. a. atrorosea* (North). NetView also more consistently recovered *E. a. abietina* and *E. a. diabolis* as distinct from each other than the PCA (Fig. 2) or sNMF (Fig. S2) analyses did.

### Evidence for recent and ongoing hybridization

There was widespread evidence for recent hybridisation within *E. abietina*. The two individuals originally suspected to have hybrid ancestry between *E. a. atrorosea* and *E. a. abietina* were consistently recovered by sNMF as sharing ancestry predominantly from clusters corresponding to *E. a. abietina* and *E. a. atrorosea* (North) (Fig. S2), and these were the only individuals recovered as F2 hybrids between these groups by NewHybrids (Fig. 4). These individuals also fell between *E. a. abietina* and *E. a. atrorosea* (North) in PCA space and in all NetView graphs. Additionally, while sNMF strongly supported the distinctness of *E. a. abietina* from *E. a. atrorosea* (North), the reverse did not apply. Only six *E. a. atrorosea* (North) individuals had <10% of their ancestry derived from the *E. a. abietina* parental population, and these consisted of the five southernmost *E. a. atrorosea* (North) samples plus one from Cecelia Forest in the central part of the *E. a. atrorosea* (North) range. The other eight *E. a. atrorosea* (North) individuals, which were not originally suspected of having mixed ancestry, had >10% of their ancestry derived from the *E. a. abietina* parental population. Of these, six individuals – including three from the northernmost sampling site at Newlands Forest and three from Cecelia Forest – were inferred by NewHybrids to be *E. a. atrorosea* (North) backcrosses, while the remaining two individuals were inferred to be non-admixed *E. a. atrorosea* (North) (Fig. 4).

**Figure 4.**
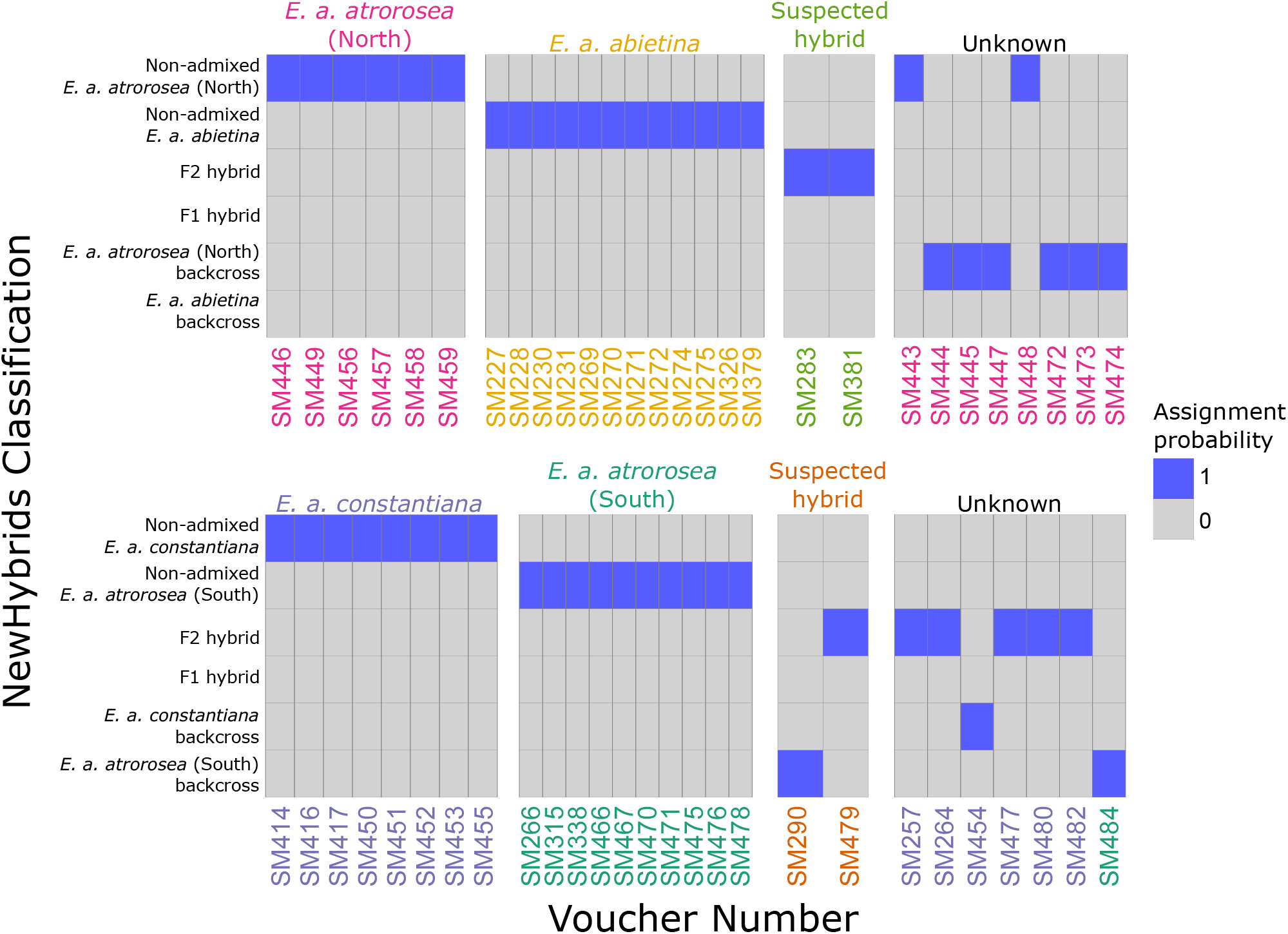
Hybrid assignment results: Posterior assignment probabilities of the various hybrid classes identifiable by NewHybrids. *Top*: *E. a. atrorosea* (North) *x E. a. abietina*; *Bottom*: *E. a. atrorosea* (South) *x E. a. constantiana*. Text colours follow Fig. 2.

The results were also complex for the analysis of hybridisation between *E. a. atrorosea* (South) and *E. a. constantiana*. According to sNMF, only one out of 11 individuals originally identified as *E. a. atrorosea* (South) had mixed ancestry, which NewHybrids recovered as an *E. a. atrorosea* (South) backcross. In contrast, six of the 14 individuals identified as *E. a. constantiana* had mixed ancestry. Five of these were recovered as F2 hybrids, including all from Blackburn Ravine and two from nearby Silvermine area just to the east, while one individual collected from the more northerly Vlakkenberg area was recovered as a *E. a. constantiana* backcross. Of the two individuals originally suspected to be hybrids between the two subspecies, one was recovered as an F2 hybrid and the other as an *E. a. atrorosea* (South) backcross.

### Phylogenetic relationships

#### All individuals included

Figure S3 depicts the results of the phylogenetic analysis with admixed individuals included. All four IQ-TREE runs returned virtually identical log-likelihood values. Overall, the tree showed successive branching forming a highly unbalanced (“ladder-like”) tree, which appeared to be significantly influenced by the presence of individuals inferred to have mixed ancestry based on sNMF and NewHybrids results. At the same time, populations identified by the analyses of population structure were readily apparent and often formed monophyletic clades. *E. a. abietina* and *E. a. diabolis* formed a well-supported clade and were confidently resolved as reciprocally monophyletic. Of the two individuals identified as *E. a. atrorosea* (North) *x E. a. abietina* F2 hybrids, one occupied a position clearly intermediate between the two populations, while the other fell within a clade containing most of the *E. a. atrorosea* (North) backcrosses and one non-admixed *E. a. atrorosea* (North) individual. The rest of the *E. a. atrorosea* (North) individuals (seven non-admixed and two backcrosses) formed a clade further towards more distant, older nodes. The individuals identified as non-admixed *E. a. constantiana* formed a well-supported clade with the inclusion of the single *E. a. constantiana* backcross individual, while all the non-admixed *E. a. atrorosea* (South) individuals not collected from Blackburn Ravine also formed a well-supported clade. A relatively poorly supported clade (bootstrap = 90%, SH-alrt = 75%) grouping between the non-admixed *E. a. atrorosea* (South) and *E. a. constantiana* clades consisted of all individuals from Blackburn Ravine regardless of prior identification. Within this clade, however, individuals identified as *E. a. atrorosea* (South) and *E. a. constantiana* each formed sub-clades that had good bootstrap support but poor SH-alrt support, and which each contained one of the individuals identified a priori as being of hybrid origin.

#### Admixed individuals excluded

Figure 5 depicts the results of the phylogenetic analysis with admixed individuals excluded. The two best IQ-TREE runs returned similar log-likelihood values (run 2: -8720997.954, run 3: -8721638.908, difference = 640.954). Most nodes of the maximum-likelihood tree received high bootstrap and SH-alrt support, particularly at deeper phylogenetic levels at which populations were distinguished. The most notable exception was the node subtending (*E. a. atrorosea* [North],(*E. a. abietina,E. a. diabolis*)), which had very low support values, meaning that the placement of *E. a. atrorosea* (North) could not be resolved. The three *E. a. atrorosea* (South) individuals from Blackburn Ravine that were not identified as being of recent hybrid origin were nevertheless recovered in an intermediate position between non-admixed *E. a. atrorosea* (South) and *E. a. constantiana*. Given that all other individuals collected from this locality were marked as putative hybrid-origin, this may indicate that these individuals have mixed ancestry of too ancient origin to have been detected by the sNMF and NewHybrids analyses.

**Figure 5.**
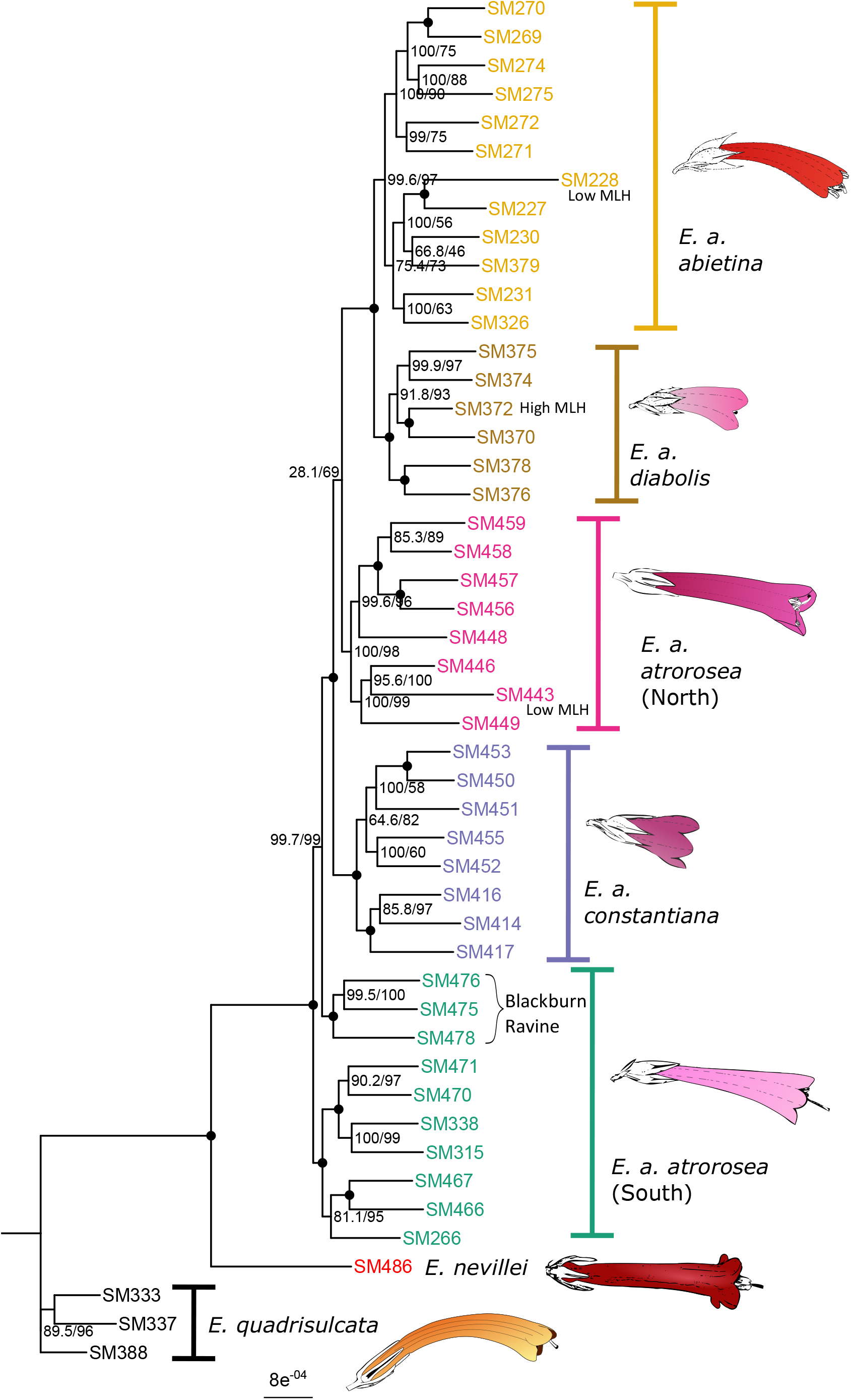
The maximum-likelihood phylogeny inferred by IQ-TREE with admixed individuals excluded. Node values indicate bootstrap/SH-alrt support, and black circles indicate full support from both measures. Text colours follow Fig. 2. Note that *E. a. atrorosea* (North) individuals form a well-supported clade but that the clade’s position in the phylogeny is unresolved. Individuals with anomalous heterozygosity and those from Blackburn Ravine are annotated.

### Population differentiation and phylogeny

The PoMo tree (Fig. 6) had the same topology as the individual-level tree with respect to population-level groupings, except that *ATRO*_*N*_ was recovered as sister to *CONS*, but with low support (bootstrap support = 72). *F*_*ST*_ was on average much higher at the species level than it was between populations within *E. abietina* (Fig. 6). Within *E. abietina*, differentiation was lowest between *ABI* and *DIAB* and highest between *CONS* and *DIAB*.

**Figure 6.**
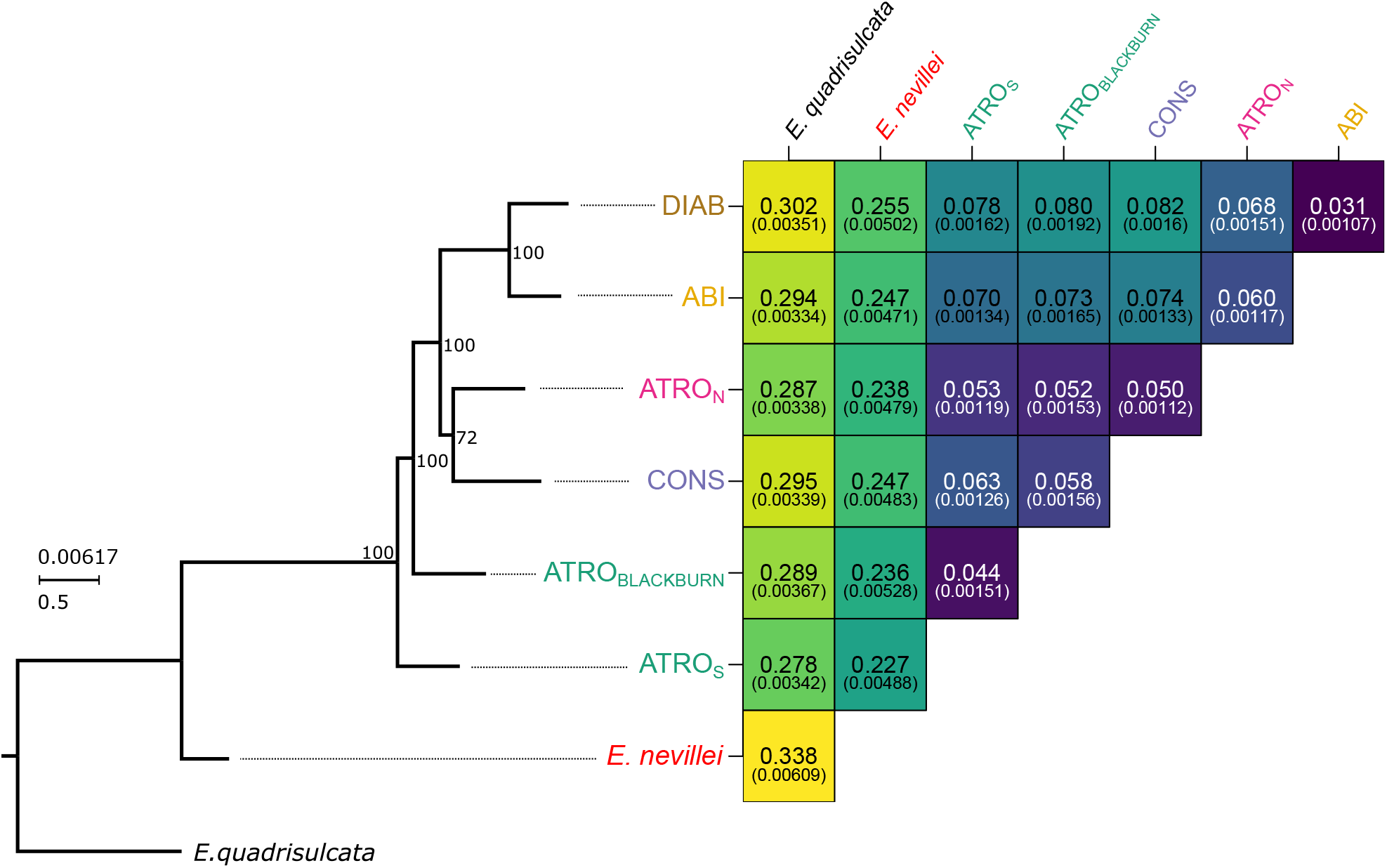
*Left*: population-level phylogeny estimated by PoMo, an incomplete lineage sorting-aware tree inference method. The tree scale shows expected substitutions per site (top) and coalescent units (bottom). Branch supports are ultrafast bootstrap. Note that the tree topology differs from that in Fig. 5 in the position of *E. a. atrorosea* (North), likely due to the exclusion from this analysis of putatively admixed individuals. *Right*: Pairwise *F*_*ST*_ between populations, with standard deviation in brackets. Darker colours indicate lower values.

### Evidence for ancient introgression

The ADMIXTOOLS2 graph search analysis results were effectively identical regardless of whether the starting tree was specified or not. When the number of allowed admixed edges (*N*_*ADMIX*_) was zero the best graph topology was identical to that of the PoMo tree in all ten runs (Fig. 7, A). With *N*_*ADMIX*_ = 1, all runs converged on the same optimal topology and essentially identical edge weights. This graph showed the ancestor of *ABI* and *DIAB* as an admixed edge with most (76%) of its ancestry derived from the ancestor of (i.e., the edge subtending) *ATRO*_*N*_ and the rest derived from the ancestor of *E. abietina* as a whole (Fig. 7, B).

**Figure 7.**
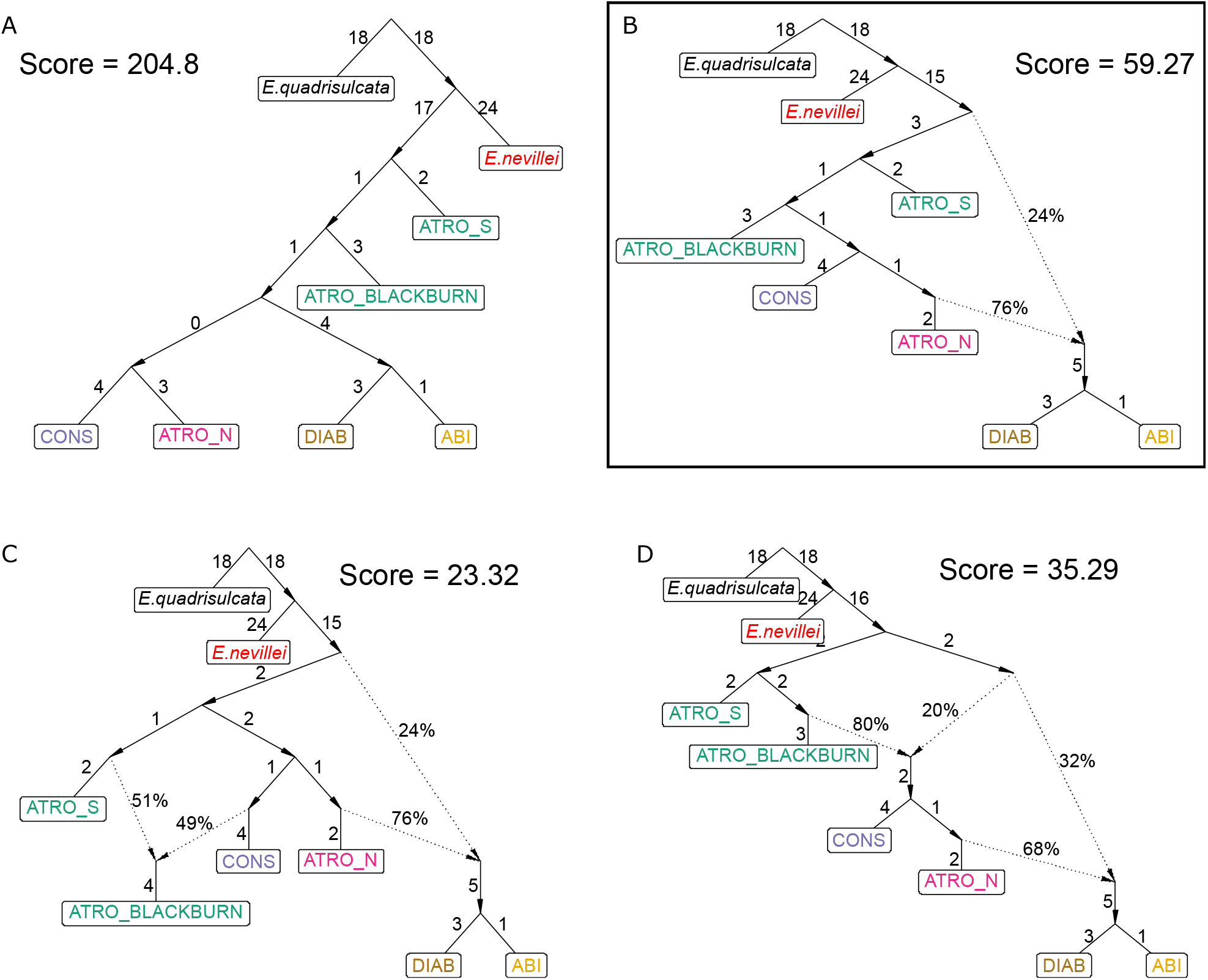
The four top-scoring admixture graphs according to ADMIXTOOLS2 for *N*_*ADMIX*_ = 0 (A), 1 (B), and 2(C, D). Numbers above solid non-admixed edges are proportional to their weight, while percentages above dashed admixture edges indicate their contribution to the admixed edge. The graph in panel B had the best performance based on its predictive power being similar to those in panels C and D while only having one admixture edge.

With *N*_*ADMIX*_ = 2, the independent runs found two distinct optimal graphs. The best-scoring graph (Fig. 7, C), found in two runs, retained the admixed edge found with *N*_*ADMIX*_ = 1 and additionally recovered *ATRO*_*BLACKBURN*_ as descending from an admixed edge with equal ancestry derived from the ancestors of *ATRO*_*S*_ and *CONS*. The next-best graph (Fig. 7, D) depicted a more complex history of admixture, with one admixed edge subtending *CONS* and *ATRO*_*N*_ deriving 80% of its ancestry from the ancestor of *ATRO*_*BLACKBURN*_ and 20% from a “ghost” lineage that was sister to the rest of *E. abietina*. This same “ghost” lineage contributed 32% of its ancestry to an admixed edge subtending *ABI* and *DIAB*, which derived the rest of its ancestry from the ancestor of *ATRO*_*N*_.

The best *N*_*ADMIX*_ = 1 graph had greater predictive power than the best *N*_*ADMIX*_ = 0 graph (mean score difference = -150.8 ± 39.5 SD, *z* = -3.82, p < 0.001). The two best *N*_*ADMIX*_ = 2 graphs had similar predictive power (mean score difference = -11.7 ±11.3 SD, *z* = -1.04, p = 0.30), however, neither had better predictive power than the best *N*_*ADMIX*_ = 1 graph (best graph: mean score difference = -31.0 ±17.7 SD, *z* = -1.75, p = 0.08; second-best graph: mean score difference = -19.3 ±15.2 SD, *z* = -1.27, p = 0.20). These results point to a single admixed edge as the most parsimonious depiction of the reticulate history of the group.

## Discussion

### Taxonomy and cryptic diversity

Given that genetic divergence between established species (e.g., between *E. abietina* and *E. nevillei*) was found to be much higher than any pairwise divergence between the various populations of *E. abietina* (Fig. 6), it is reasonable to conclude that *E. abietina* constitutes a single – if highly variable – species. On the other hand, there is mixed support for the current subspecific treatment of the complex. In particular, what is currently recognised as *E. a. atrorosea* appears to consist of two non-sister lineages that are geographically isolated but phenotypically very similar. Such “cryptic” diversity has come to be detected frequently in a range of organisms (Bickford et al., 2007), particularly since the advent of Next Generation Sequencing-based genotyping in non-model organisms (e.g., Hinojosa et al., 2019; Blair et al., 2019; Lutsak, 2020; Boucher et al., 2021; Daïnou et al., 2016). The *E. a. atrorosea* (South) lineage appears to be widespread on sandstone-derived soils south of Table Mountain, whereas *E. a. atrorosea* (North) is seemingly restricted to the granite- and shale-derived soils of the eastern slopes of Table Mountain and Constantiaberg (see Fig. 1). None of our results suggested current or ancient hybridization between these two lineages, lending support to their independence.

### Floral trait evolution

Despite the close relationships and weak reproductive barriers between the populations of *E. abietina*, shifts between pollination syndromes (as approximated by flower tube length) occurred at least twice in the complex. Given the phylogenetic results, the most parsimonious reconstruction of floral trait evolution is that long tubes are plesiomorphic and short tubes were derived in *E. a. constantiana* and *E. a. diabolis* independently. Such a pattern, combined with incomplete reproductive isolation between sympatric short-and long-tubed populations (in the case of *E. a. constantiana* and *E. a. atrorosea* [South]), implies that strong selective forces drive and maintain shifts to insect pollination.

What might drive shifts to different modes of pollination? At least for *E. a. constantiana*, the Cape Honeybee (*Apis mellifera* subsp. *capensis*) seems likely to be the primary pollinator based on numerous personal observations (e.g., https://www.inaturalist.org/observations/26399879, https://www.inaturalist.org/observations/131065480). Shifting from bird to bee pollination could be interpreted as a shift to a less specialised pollination system (Cronk and Ojeda, 2008). Johnson (1996), for example, suggested that sunbirds are more reliable pollinators than insects and that this, in addition to the relatively low abundance of insects in fynbos, especially at higher elevations, drove frequent shifts to sunbird pollination in many CFR lineages. However, van der Niet et al. (2020) showed that individual honeybees visiting two co-occurring Cape *Erica* species exhibited remarkable floral constancy, in that during foraging bouts they tended to consistently prefer one or the other species rather than visiting both. Such constancy presumably explained the authors’ finding of extremely low rates of interspecific pollen transfer. Based on a detailed analysis of a community of co-occurring insect-pollinated *Erica*, Bouman et al. (2017) suggested that, rather than incurring a fitness cost, bee pollination may instead benefit co-occurring species by enabling them to collectively attract pollinators while simultaneously avoiding cross-species pollen flow. Such benefits may explain the independent evolution of short-tubed flowers in *E. a. constantiana* and *E. a. diabolis*. Overall, our results highlight the variability of selection on floral traits and point to *Erica* as a whole being highly sensitive to such selection.

Despite the evident lability of floral traits in *Erica*, their link to the genus’s diversification is less certain. Our results showed that pollination syndrome differentiation cannot be assumed to be a reliable indicator of reproductive isolation. In the CFR, floral tube length variation has been shown to cause intraspecific reproductive isolation in *Erica* (Newman and Johnson, 2021) and other angiosperm groups (e.g., Minnaar et al., 2019). However, a global-scale meta-analysis showed that floral trait divergence on its own is a poor predictor of speciation, and that it instead almost always acts in concert with other factors – such as geographic and habitat isolation – to effect speciation (Kay and Sargent, 2009). Our results support this assertion in that all but one individual showing mixed ancestry between *E. a. constantiana* and *E. a. atrorosea* (South) occurred at the same locality (Blackburn Ravine), whereas putatively non-admixed individuals occurred in areas where only short- or long-tubed individuals, but not both, are present. We therefore suggest that floral trait divergence between *E. a. constantiana* and *E. a. atrorosea* (South) and between *E. a. diabolis* and *E. a. abietina* resulted from a combination of pollination-driven selection and geographic isolation resulting in genetic drift. The cryptic diversity within *E. a. atrorosea* – which we suggest was driven by geographic isolation coupled with adaptation of the northern lineage to relatively nutrient-rich soils – further illustrates that divergence can occur without a pollinator shift.

### Hybridization and introgression

Although *E. abietina* can be grouped into genetically distinct, geographically separated and phenotypically recognisable units, these cannot be said to represent completely isolated lineages. The discovery of several putative hybrids of recent origin – which occurred at localities where more than one subspecies was present or where their ranges met (Fig. 1, 4) – shows that gene flow between certain populations is ongoing and probably frequent. Interspecific hybridisation is well-known in Cape *Erica* (Oliver and Oliver, 2002, 2005), and does not always involve closely related species (e.g., Oliver, 1986). Interestingly, in this study none of the putative hybrid individuals was inferred to be first-generation and most were backcrosses. This implies that hybrids are fertile and liable to generate “hybrid swarms”. Such a hybrid swarm may exist at Blackburn Ravine where *E. a. atrorosea* (South) and *E. a. constantiana* appear to have formed a population with a large proportion of admixed individuals capable of backcrossing with either parental lineage.

In contrast, recent introgression between *E. a. abietina* and *E. a. atrorosea* (North) seems to have been unidirectional. This was particularly evident from the sNMF analyses, in which all *E. a. abietina* individuals were inferred to be genetically non-admixed, whereas *E. a. atrorosea* (North) individuals showed a pattern of increasingly mixed ancestry from south to north. This could reflect a case of recent secondary contact between these lineages; however, this would imply recent range expansion in one or both of these populations. This seems unlikely given that genetic diversity (measured by heterozygosity) did not show a clear relationship with geography in either of them (Fig. S1). Nevertheless, denser geographic sampling of both populations enabling formal analyses aimed at detecting range expansions (e.g., He et al., 2017; Peter and Slatkin, 2013) would be a valuable endeavour. Alternatively, the pattern may instead reflect long-standing gene flow from *E. a. abietina* into *E. a. atrorosea* (North) that has not been sufficient to cause gene swamping (Lenormand, 2002; Bridle and Vines, 2007). *Erica a. abietina* is likely to be adapted to extremely nutrient-poor sandstone-derived soils such as exist on Table Mountain’s upper plateau (Compton, 2004), and such specialisation has been shown to limit the ability of fynbos plants to adapt to more nutrient-rich soils (Verboom et al., 2017). This may mean that *E. a. abietina* alleles are maladaptive for *E. a. abietina x E. a. atrorosea* (North) hybrids occupying the lower eastern slopes – whose soils are relatively rich in nutrients (Compton, 2004; Cramer et al., 2018, Fig. 1) – resulting in what has been termed “migration-selection equilibrium” (Lenormand, 2002). Studies investigating the factors that determine the geographic range limits of these populations (e.g., local adaptation, competition, dispersal limitation; Gaston, 2009) would help to differentiate these possibilities.

Evidence of present-day hybridisation does not necessarily mean that introgression has influenced diversification (e.g., Kessler et al., 2022; Jordan et al., 2017; Westbury et al., 2019), however, our analysis based on *f3* -statistics provides good evidence that *E. abietina* does have a reticulate evolutionary history. Perhaps the most biologically sensible interpretation of the best-fitting admixture graph (Fig. 7, B) is that *E. a. abietina*/*E. a. diabolis* split from the rest of the complex while simultaneously deriving some (roughly one quarter) of their genetic makeup from a more distant relative ancestral to or close to *E. nevillei*. This interpretation implies the influence of a “ghost” lineage: one that is unknown, unsampled or extinct and which interbred historically with an extant lineage (Ottenburghs, 2020). Ghost introgression appears to be common across the tree of life (e.g., Meyer et al., 2012; Green et al., 2010; Barlow et al., 2018; Maier et al., 2022) and may be an important driver of diversification in rapidly evolving lineages (Ottenburghs, 2020). *Erica nevillei* and *E. quadrisulcata* are both narrow-range endemics confined to rocky outcrops at relatively high elevations (Oliver and Oliver, 2002). Given this, it is surprising that the seemingly suitable upper plateau of Table Mountain does not host these or a related species with a similar niche. However, assuming that this absence is the result of extinction could explain why ghost introgression is inferred. Such speculation could be tested with improved sampling of the relevant populations and a more thorough genotyping method (such as whole-genome sequencing) which would allow for more detailed and powerful analyses (e.g., Mondal et al., 2019), including those that can detect adaptive introgression of genomic regions (e.g., Racimo et al., 2015).

## Conclusions

This study adds to the small but growing and much-needed body of literature focused on understanding speciation in action in the Fynbos flora (Ellis et al., 2014; Barraclough, 2006; Prunier and Holsinger, 2010; Prunier et al., 2017; Lexer et al., 2014). We find that the subspecific classification of *E. abietina* is only a partial reflection of its evolutionary history. Notably, there is evidence of cryptic diversity in the complex, which may have gone unrecognised due to a focus on floral traits in the taxonomic literature (Oliver and Oliver, 2002). We also find that floral trait divergence is likely to be driven by strong selective forces but that it is unlikely to drive lineage divergence without the action of additional factors, particularly geographic isolation. Finally, apart from ongoing gene flow in the complex revealed by the presence of subspecific hybrids, we find evidence that ancient gene flow may have also influenced the complex’s evolution in the form of “ghost” introgression from a now-extinct lineage. Exploring the possibility of links between introgression and floral trait evolution in *Erica* may provide important insights into the spectacular diversification of the genus (see e.g., Nelson et al., 2021). Overall our results paint a picture of a highly dynamic system whose evolutionary history has been shaped by diverse, interacting forces (Donoghue and Sanderson, 2015). Refining this picture will undoubtedly further our understanding of diversification in *Erica* and beyond.

## Acknowledgements

Primary funding for the project was provided by the Deutsche Forschungsgemeinschaft (PI 1169/1-2). The Genomics Core Facility (GCF) at the University of Bergen, which is a part of the NorSeq consortium, provided services on whole-genome sequencing of *Erica cerinthoides*; GCF is supported in part by major grants from the Research Council of Norway (grant no. 245979/F50), Bergen Research Foundation (BFS) (grant no. BFS2017TMT04 and BFS2017TMT08), and Trond Mohn Foundation (TMS). Computing facilities were provided by the University of Cape Town’s ICTS High Performance Computing team (hpc.uct.ac.za) and the Centre for High Performance Computing at the University of Bayreuth (bzhpc.uni-bayreuth.de). Collections were made under permits from CapeNature (CN35-31-8281) and SANParks (CRC/2019-2020/004–2019/V1). Voucher specimens were deposited at the Compton Herbarium (NBG).

## Author contributions

NN and SM conceived the study. SM collected samples, did the lab work, and analysed the data. SM wrote the initial draft with input by MP and NN. All authors reviewed the final manuscript.

## Data availability

Raw reads will be deposited in GenBank and other associated data including genotypic data, metadata and code will be made available upon publication.

## Supporting Information

**Figure S1**. Multilocus heterozygosity (MLH) of all individuals in the study.

**Figure S2**. Mean individual ancestry proportions estimated by sNMF.

**Figure S3**. Maximum-likelihood phylogeny inferred by IQ-TREE with all individuals included.

**Figure S4**. Read redundancy comparison between sequencing batches.

**Table S1**. Table summarising variant filtering procedures and features of the genotype matrices.

**Table S2**. Voucher information.

