## Supporting Information (SI) for "Geographic isolation enables recurrent pollinator shifts despite hybridisation in the Cape’s hyperdiverse heathers" (from another version of this preprint: version 1, 2023-07-19)

for

The dynamics of species divergence in South Africa's megadiverse Cape region.

#### 1: Supporting Figures

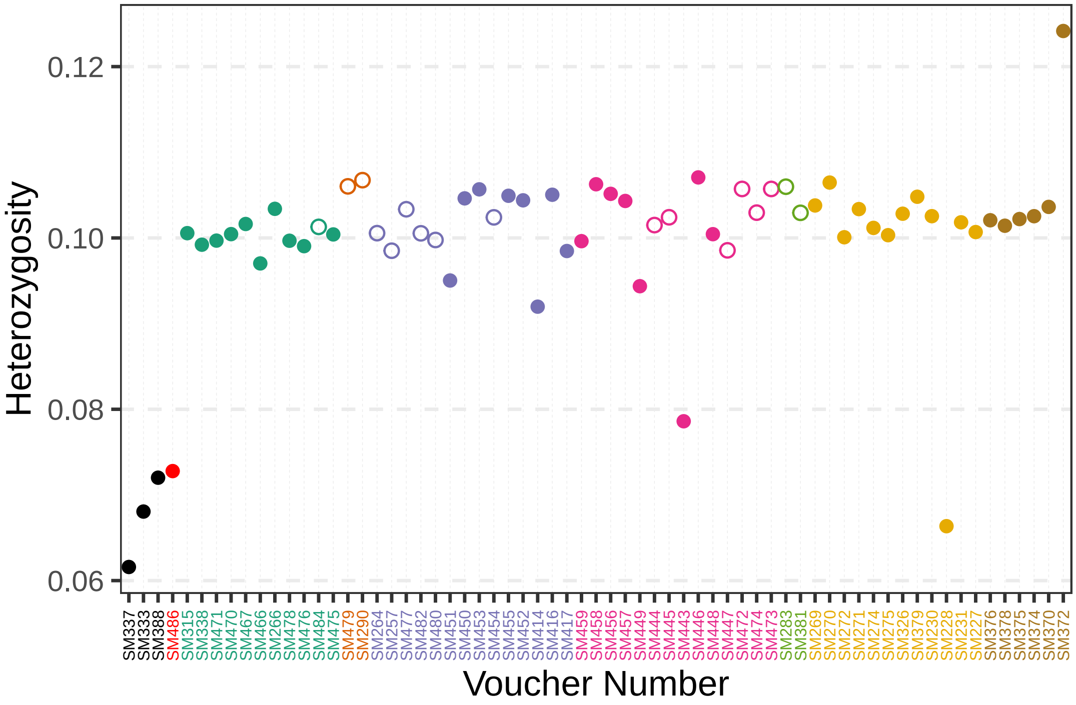


**Figure S1** Multilocus heterozygosity (MLH) of all individuals in the study. Text and point colours follow Fig. 2 in the main text. Unfilled dots indicate individuals identified as having putative hybrid origin based on the NewHybrids analyses. Within groups, individuals are arranged by latitude from south to north.

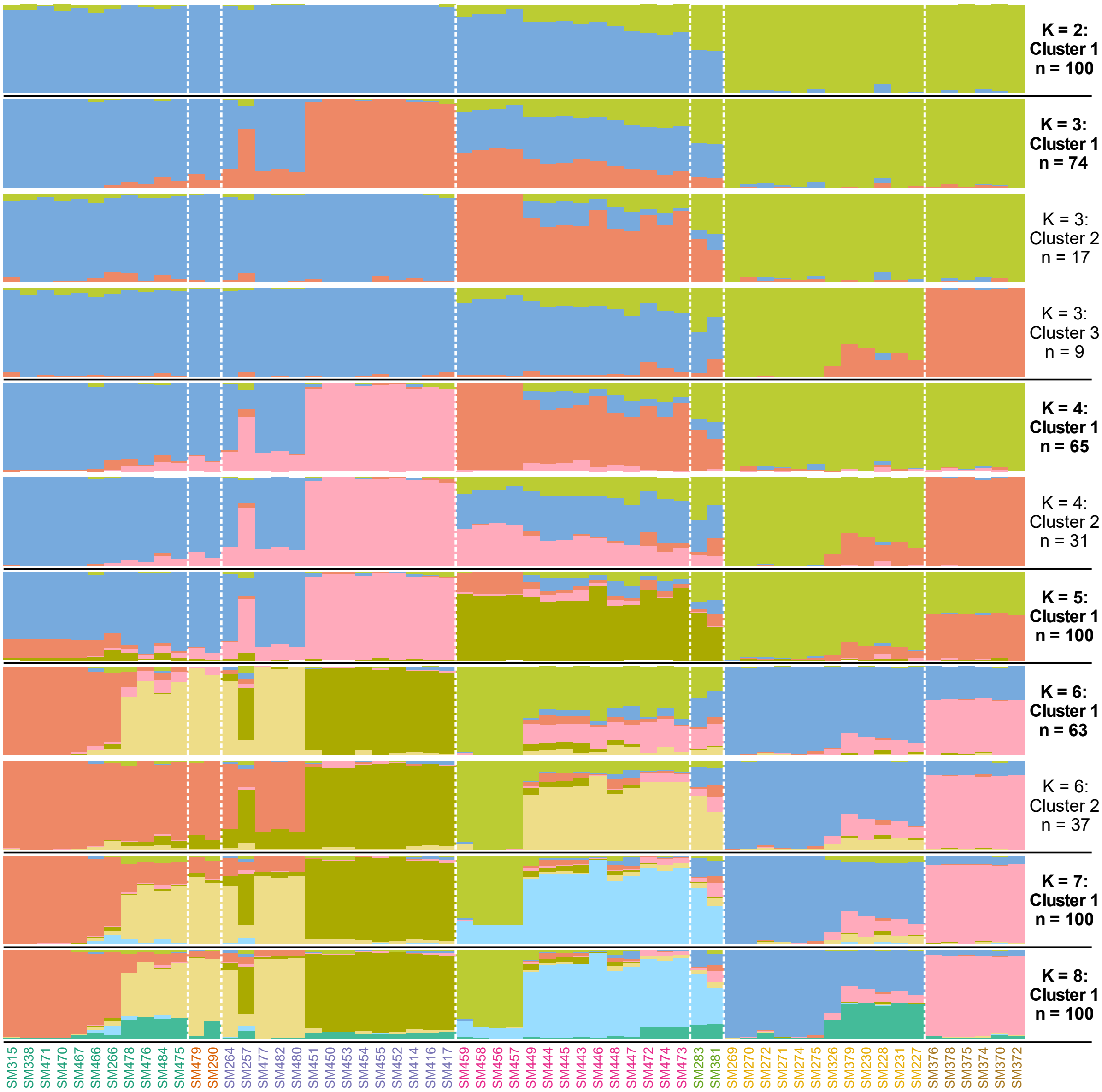


**Figure S2** Mean individual ancestry proportions estimated by sNMF for $K = 2$–8 with 100 runs for each $K$ clustered by the CLUMPAK method. One cluster for $K = 3$ with $n = 4$ runs is not shown. Voucher numbers are shown below individuals., with text colours following Fig. 2 in the main text. Within groups, individuals are arranged by latitude from south to north. Note especially the presence of more than one potentially optimal solution for values of $K = 3$, 4, and 6; the increasingly mixed ancestry in *E. a. atrorosea* (North) from south to north; and the ability of the method to distinguish *E. a. diabolis* from *E. a. abietina* in many runs.

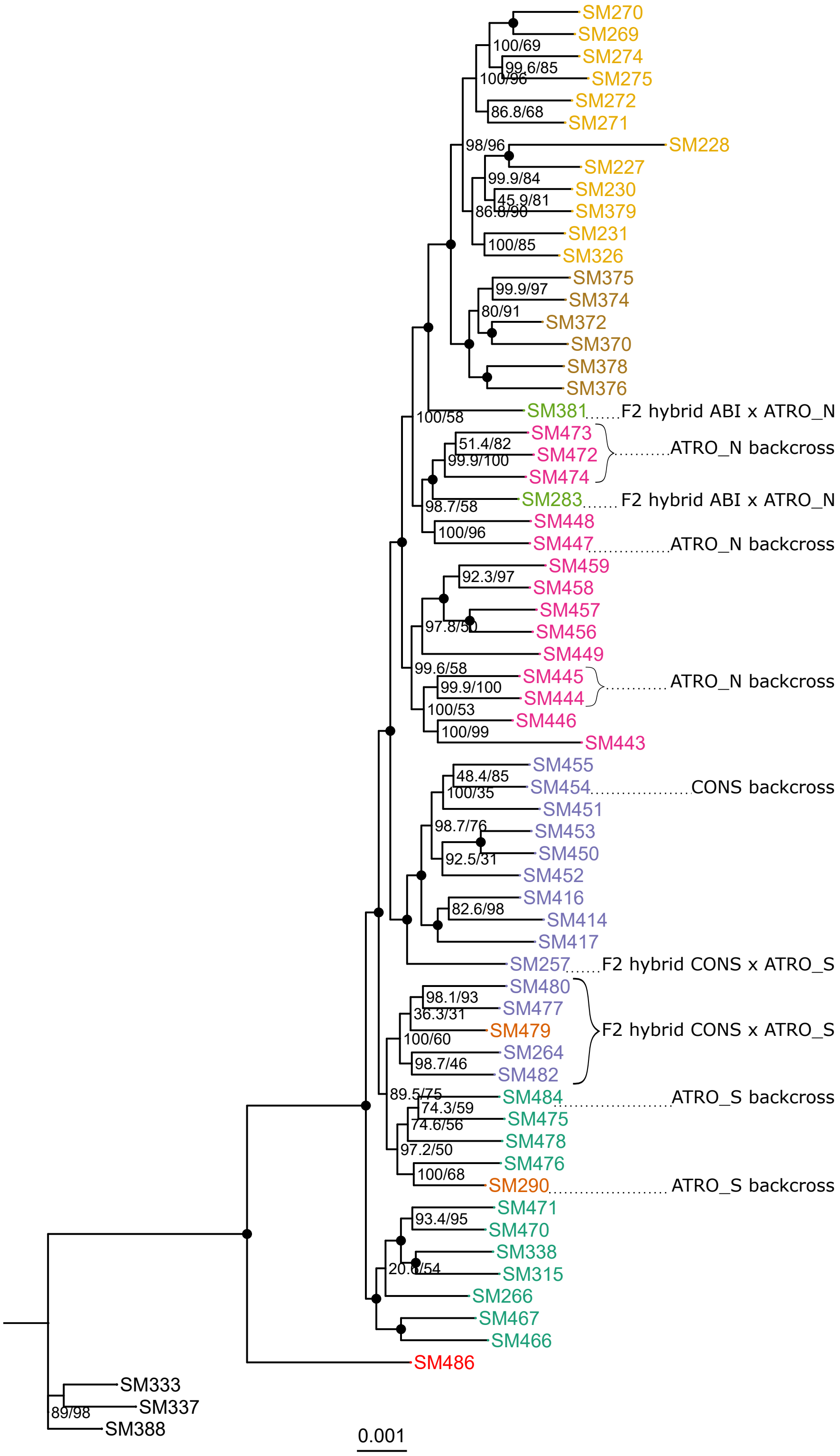


**Figure S3** The maximum-likelihood phylogeny inferred by IQ-TREE with all individuals included. Recent hybrids detected by NewHybrids are annotated. Node values indicate bootstrap/SH-alrt support, and black circles indicate full support from both measures. Text colours follow Fig. 2 in the main text.

### 2: Supporting Tables

**Table S1** Table summarising the variant filtering applied and resulting features of the genotype matrices.

| Set name | Filtering criteria (sites kept) | Total sites | Contigs | SNPs | Invariant sites | Genotyping rate |
|---|---|---|---|---|---|---|
| *Variant sites only* | | | | | | |
| VAR_RAW | All variant types; Read mapping quality $\geq$ 10. | 7 800 988 | 186 414 | 6 373 677 | - | 0.43 |
| SNP_HQ | Decompose complex variants; Biallelic SNPs only; GT DP $\geq$ 3; Missingness $\leq$ 50%; MAF $\geq$ 0.01; AB = 0 OR 0.25 ¡ AB ¡ 0.75; 0.9 ¡ MQM/MQMR ¡ 1.1. | 771 049 | 45 804 | 771 049 | - | 0.81 |
| SNP_HQ_ExcessHet | Excess heterozygosity: p-value ¿ 0.2. | 738 178 | 45 609 | 738 178 | - | 0.81 |
| SNP_m10 | Missingness $\leq$ 10%. | 292 849 | 20 697 | 292 849 | - | 0.96 |
| SNP_m10_LD | Linkage disequilibrium. | 179 927 | 20 695 | 179 927 | - | 0.96 |
| SNP_m10_LD_maf04 | MAF $\geq$ 0.04. | 94 883 | 18 969 | 94 883 | - | 0.96 |
| *Variant plus invariant sites* | | | | | | |
| ALL_m10 | Missingness $\leq$ 10%. | 4 993 526 | 22 920 | 292 849 | 4 700 677 | 0.97 |

GT: genotype; DP: read depth; MAF: minor allele frequency; AB: allele balance;
MQM(R): mean mapping quality of reads supporting alternate (reference) allele

**Table S2** Voucher information. In bold is the voucher number of the individual sequenced in both sequencing batches (SM484).

| Voucher No. | Sample No. | Batch | Latitude | Longitude | Taxon | Genetic group | iNaturalist ID |
|---|---|---|---|---|---|---|---|
| SM337 | S166z | 2 | -34.25393600 | 18.46956497 | *Erica quadrisulcata* | *E. quadrisulcata* | 32097433 |
| SM333 | S214z | 2 | -34.23876297 | 18.46307300 | *Erica quadrisulcata* | *E. quadrisulcata* | 32097429 |
| SM388 | S279B | 2 | -34.21381006 | 18.45148847 | *Erica quadrisulcata* | *E. quadrisulcata* | 37615208 |
| SM486 | S378 | 1 | -34.05089909 | 18.36669613 | *Erica nevillei* | *E. nevillei* | 63581343 |
| SM315 | S142 | 2 | -34.31851300 | 18.41953600 | *E. a. atrorosea* | *E. a. atrorosea* (South) | 30927503 |
| SM338 | S167 | 2 | -34.26035300 | 18.46235797 | *E. a. atrorosea* | *E. a. atrorosea* (South) | 32097434 |
| SM471 | S331 | 2 | -34.18790687 | 18.37421503 | *E. a. atrorosea* | *E. a. atrorosea* (South) | 62543624 |
| SM470 | S330 | 2 | -34.18193497 | 18.37504584 | *E. a. atrorosea* | *E. a. atrorosea* (South) | 62543568 |
| SM467 | S354 | 2 | -34.09040187 | 18.42164788 | *E. a. atrorosea* | *E. a. atrorosea* (South) | 62543342 |
| SM466 | S353B | 1 | -34.09039687 | 18.42165895 | *E. a. atrorosea* | *E. a. atrorosea* (South) | 62543295 |
| SM266 | S197 | 2 | -34.07493800 | 18.39962600 | *E. a. atrorosea* | *E. a. atrorosea* (South) | 26399907 |
| SM478 | S370 | 2 | -34.05868184 | 18.37431192 | *E. a. atrorosea* | *E. a. atrorosea* (South) | 63160143 |
| SM476 | S368z | 2 | -34.05853602 | 18.37298490 | *E. a. atrorosea* | *E. a. atrorosea* (South) | 63159779 |
| **SM484** | S376 | 1 | -34.05843102 | 18.37782595 | *E. a. atrorosea* | *E. a. atrorosea* (South) | 63163263 |
| **SM484** | S376D | 2 | -34.05843102 | 18.37782595 | *E. a. atrorosea* | *E. a. atrorosea* (South) | 63163263 |
| SM475 | S366 | 2 | -34.05658386 | 18.37230496 | *E. a. atrorosea* | *E. a. atrorosea* (South) | 63159621 |
| SM479 | S371 | 2 | -34.05859102 | 18.37438401 | *E. a. constantiana x E. a. atrorosea* | *E. a. constantiana x E. a. atrorosea* (South) | 63162993 |
| SM290 | S367 | 2 | -34.05852000 | 18.37296500 | *E. a. constantiana x E. a. atrorosea* | *E. a. constantiana x E. a. atrorosea* (South) | 29499592 |
| SM264 | S92zA | 2 | -34.06371900 | 18.38555600 | *E. a. constantiana* | *E. a. constantiana* | 26399903 |
| SM257 | S110zA | 2 | -34.06253000 | 18.39633600 | *E. a. constantiana* | *E. a. constantiana* | 26399851 |
| SM477 | S369 | 2 | -34.05895711 | 18.37424889 | *E. a. constantiana* | *E. a. constantiana* | 63159964 |
| SM482 | S374 | 2 | -34.05861101 | 18.37419994 | *E. a. constantiana* | *E. a. constantiana* | 63160382 |
| SM480 | S372 | 2 | -34.05859102 | 18.37438401 | *E. a. constantiana* | *E. a. constantiana* | 63160250 |
| SM451 | S337 | 2 | -34.02258007 | 18.40146594 | *E. a. constantiana* | *E. a. constantiana* | 57652605 |
| SM450 | S336 | 2 | -34.02241807 | 18.40491090 | *E. a. constantiana* | *E. a. constantiana* | 57651903 |
| SM453 | S338 | 2 | -34.02234387 | 18.40407606 | *E. a. constantiana* | *E. a. constantiana* | 57653130 |
| SM454 | S340 | 2 | -34.02230691 | 18.40522405 | *E. a. constantiana* | *E. a. constantiana* | 57653495 |
| SM455 | S341 | 2 | -34.02205710 | 18.40527602 | *E. a. constantiana* | *E. a. constantiana* | 57654128 |
| SM452 | S339 | 2 | -34.02176587 | 18.40137105 | *E. a. constantiana* | *E. a. constantiana* | 57652774 |
| SM414 | S314 | 2 | -33.99948100 | 18.40007991 | *E. a. constantiana* | *E. a. constantiana* | 40627133 |
| SM416 | S316 | 2 | -33.99816791 | 18.40062842 | *E. a. constantiana* | *E. a. constantiana* | 40647963 |
| SM417 | S317 | 1 | -33.99532737 | 18.40196483 | *E. a. constantiana* | *E. a. constantiana* | 40648239 |
| SM459 | S348 | 2 | -34.02265593 | 18.40658594 | *E. a. atrorosea* | *E. a. atrorosea* (North) | 58057956 |
| SM458 | S347 | 2 | -34.02245503 | 18.40642802 | *E. a. atrorosea* | *E. a. atrorosea* (North) | 58057565 |
| SM456 | S342 | 2 | -34.02231108 | 18.40626206 | *E. a. atrorosea* | *E. a. atrorosea* (North) | 57753114 |
| SM457 | S343 | 2 | -34.02225300 | 18.40622485 | *E. a. atrorosea* | *E. a. atrorosea* (North) | 58057054 |
| SM449 | S335 | 2 | -34.01266297 | 18.41973715 | *E. a. atrorosea* | *E. a. atrorosea* (North) | 56335274 |
| SM444 | S326 | 1 | -34.00541992 | 18.41663785 | *E. a. atrorosea* | *E. a. atrorosea* (North) | 55685470 |
| SM445 | S327 | 1 | -34.00364943 | 18.41361299 | *E. a. atrorosea* | *E. a. atrorosea* (North) | 56165232 |
| SM443 | S325 | 2 | -34.00298209 | 18.42126902 | *E. a. atrorosea* | *E. a. atrorosea* (North) | 55685271 |
| SM446 | S332 | 2 | -34.00104589 | 18.41553614 | *E. a. atrorosea* | *E. a. atrorosea* (North) | 56214476 |
| SM448 | S334 | 1 | -33.99812511 | 18.42417587 | *E. a. atrorosea* | *E. a. atrorosea* (North) | 56334836 |
| SM447 | S333 | 1 | -33.99289091 | 18.42598803 | *E. a. atrorosea* | *E. a. atrorosea* (North) | 56214830 |
| SM472 | S344 | 2 | -33.97574005 | 18.44338987 | *E. a. atrorosea* | *E. a. atrorosea* (North) | 62697322 |
| SM474 | S346 | 2 | -33.97562800 | 18.44313707 | *E. a. atrorosea* | *E. a. atrorosea* (North) | 63159540 |
| SM473 | S345 | 2 | -33.97562494 | 18.44316792 | *E. a. atrorosea* | *E. a. atrorosea* (North) | 63159457 |
| SM283 | S120 | 2 | -33.98199100 | 18.43313500 | *E. a. abietina x E. a. atrorosea* | *E. a. abietina x E. a. atrorosea* (North) | 28933719 |
| SM381 | S191 | 2 | -33.96038300 | 18.44319197 | *E. a. abietina x E. a. atrorosea* | *E. a. abietina x E. a. atrorosea* (North) | 37482868 |
| SM269 | S384 | 2 | -33.98950000 | 18.41341600 | *E. a. abietina* | *E. a. abietina* | 28425132 |
| SM270 | S385 | 1 | -33.98942100 | 18.41317000 | *E. a. abietina* | *E. a. abietina* | 28425133 |
| SM272 | S387 | 1 | -33.98891100 | 18.41154097 | *E. a. abietina* | *E. a. abietina* | 28425139 |
| SM271 | S386 | 2 | -33.98888997 | 18.41293500 | *E. a. abietina* | *E. a. abietina* | 28425137 |
| SM274 | S388 | 2 | -33.98807800 | 18.41380200 | *E. a. abietina* | *E. a. abietina* | 28425141 |
| SM275 | S249 | 2 | -33.98794697 | 18.41441700 | *E. a. abietina* | *E. a. abietina* | 28425144 |
| SM326 | S160 | 2 | -33.96921497 | 18.38995997 | *E. a. abietina* | *E. a. abietina* | 30927835 |
| SM379 | S189 | 2 | -33.95646097 | 18.43214497 | *E. a. abietina* | *E. a. abietina* | 37482865 |
| SM230 | S226zA | 2 | -33.95631900 | 18.42578100 | *E. a. abietina* | *E. a. abietina* | 25265906 |
| SM228 | S380 | 1 | -33.95580500 | 18.42421424 | *E. a. abietina* | *E. a. abietina* | 25265904 |
| SM231 | S382 | 2 | -33.95577800 | 18.42717300 | *E. a. abietina* | *E. a. abietina* | 25265913 |
| SM227 | S379B | 1 | -33.95529900 | 18.42420300 | *E. a. abietina* | *E. a. abietina* | 25265963 |
| SM376 | S186 | 2 | -33.95436300 | 18.43880800 | *E. a. diabolis* | *E. a. diabolis* | 37482861 |
| SM378 | S188 | 2 | -33.95403797 | 18.43765397 | *E. a. diabolis* | *E. a. diabolis* | 37482864 |
| SM375 | S185 | 2 | -33.95335597 | 18.43986597 | *E. a. diabolis* | *E. a. diabolis* | 37482860 |
| SM374 | S184 | 2 | -33.95321597 | 18.43978200 | *E. a. diabolis* | *E. a. diabolis* | 37482859 |
| SM370 | S180 | 2 | -33.95213320 | 18.44620116 | *E. a. diabolis* | *E. a. diabolis* | 37011954 |
| SM372 | S275 | 2 | -33.95190820 | 18.44614618 | *E. a. diabolis* | *E. a. diabolis* | 37291782 |

iNaturalist observations can be viewed at `inaturalist.org/observations/<iNaturalistID>`.

#### 3: Assessing the impact of lower sequencing effort

Using a lower sequencing effort is more cost effective but could compromise data quality by reducing the accuracy of genotype calls and increasing the rate of missing data. The first batch consisted of 12 samples sequenced to an estimated 120 Mbp each, while the second batch consisted of 54 samples sequenced to an estimated 240 Mbp each (i.e. batch 2 had much greater sequencing effort). To evaluate whether the libraries were sequenced sufficiently to cover most of their complexity, we estimated redundancy in the read pool of each sample using *bbcountunique.sh* from BBTools v. 38.90 (BBMap - Bushnell B. - `sourceforge.net/projects/bbmap/`) with default parameter values. In this analysis read pairs are inspected in random chunks of 25 000, and for each chunk the proportion of read pairs (including all previous chunks) that are completely unique is estimated. A well-sequenced library should show a rapid decline in the proportion of unique reads, tending towards zero as more reads are inspected. The first and second batches of sequencing resulted, respectively, in a mean of 0.942 ($\pm$0.168 SD) and 1.96 ($\pm$0.395 SD) million read pairs per sample. Mean GC content across all samples was 39.0% ($\pm$0.349% SD) and did not differ between batches (linear model, $F(1,64) = 1.564$, $p = 0.216$). The read redundancy analysis showed that batch 2 had more unique reads than batch 1, but also that batch 2 showed somewhat diminishing returns as the number of unique reads encountered began to plateau beyond *ca.* 1.5 million read pairs (Fig. S4). The shape of the curves, with the proportion of unique reads encountered dropping rapidly as the number of reads inspected grew, suggested that for both batches sequencing effort was sufficient to capture most of the library complexity.

To test the effect of sequencing effort on genotype calling accuracy, SNPs were called separately for the read sets from the sample that was sequenced twice (once in each batch) using Freebayes v. 1.3.4 (Garrison and Marth, 2012) and the rate of discordance in genotype calls was estimated. Mean genotype depth (number of reads for called genotypes) was 16.9 in batch 1 and 31.4 in batch 2. Of the 29 998 sites (including all variant types) that were called as heterozygous based on batch 2 reads, 235 were called as homozygous and 185 were called as other genotypes based on batch 1 reads; therefore, assuming the batch 2 calls to be correct gives an error rate of 1.4% induced by lower sequencing effort. When including only SNPs, this putative error rate dropped to just 0.82% (of the 24 753 heterozygous SNPs, 205 were called as homozygous). Given the results of these analyses, we concluded that there was no significant batch effect and that genotype calls for batch 1 individuals would be accurate despite their lower depth.

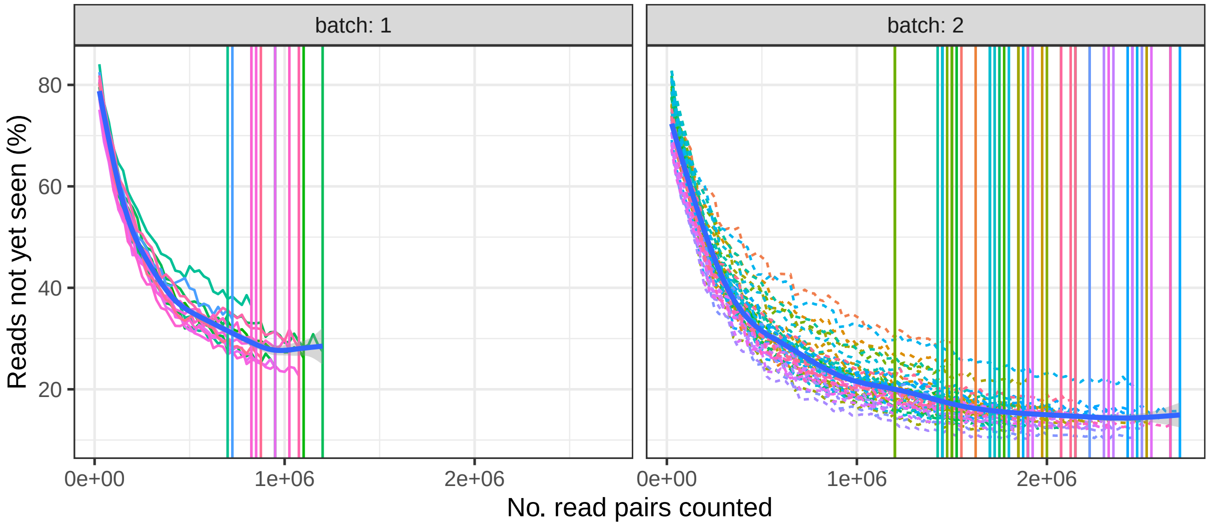


**Figure S4** Redundancy analysis results from *bbcountunique.sh*. Read pairs are inspected in chunks of 25 000, and for each chunk the proportion of read pairs (including all previous chunks) that are completely unique is estimated. Colours are to aid in distinguishing estimates from different individuals. The plots show that unique reads quickly become the minority, meaning that the libraries were sequenced sufficiently in both batches.

### 4: DNA extraction protocol for Erica leaf material

#### Note

This protocol was adapted from that of Inglis et al. (2018). It works best on young leaf material (in *Erica* the youngest leaves are located at the branch tips). If using fresh material, it is best to grind the leaves using liquid N, but they can also be ground in SWB (see below) in 2 mL Eppendorf tubes with steel beads using a TissueLyser or a similar product.

The primary modifications to the Inglis et al. (2018) protocol are as follows:

1. Instead of CTAB, SDS is used as the detergent for cell lysis. This allows for a purification step in which K Acetate is added after lysis, causing proteins and polysaccharides to precipitate along with SDS during a cooling step.
2. A combination of NaCl and Na Acetate is used to inhibit the co-precipitation of polysaccharides with DNA after the addition of isopropanol. NaCl increases the solubility of polysaccharides but not DNA, but can also cause the isopropanol to come out of solution. By experimentation, the addition of Na Acetate was found to prevent the latter from occurring.

#### Materials

- SWB: Sorbitol wash buffer (2,000 µL per sample): 100 mM Tris-HCl pH 8.0, 0.35 M Sorbitol, 1% PVP, 10 mM EDTA pH 8.0. **STORE AT 4°C** for up to 6 months.
- 20% SDS (Sodium dodecyl sulphate) (80 µL per sample)
- Extraction buffer (800 µL per sample): 500mM NaCl, 100mMTris-HCl, 50mM EDTA, adjusted to pH 8.0.
- RNase A: 100 mg/mL (Qiagen Cat. No. / ID: 19101)
- 5M K Acetate (KAc; **STORE AT 4°C.**) (80 µL per sample)
- CIA: Chloroform:Isoamyl alcohol (24:1) (600 µL per sample).
- 5M NaCl: (150 µL per sample).
- 3M Na Acetate, pH 5.2 (50 µL per sample).
- Eppendorf tubes: 1 x 2 mL tube, 2 x 1.5 mL tubes.
- Isopropanol: 500 µL per sample.
- Ethanol (70%): 1000 µL per sample.
- TE elution buffer: 105 µL per sample.

#### Protocol

1. Prepare a working solution of Extraction buffer, adding 2 µL RNase A per sample and 1% $\beta$-mercaptoethanol.
2. Prepare a working solution of SWB, adding 1% $\beta$-mercaptoethanol.
3. Prepare water bath or oven at 65°C.
4. Weigh out 25-35 mg dry leaf material per sample.

5. Add dry leaf material to 2 mL Eppendorf tube with at least two grinding beads. I used two 2.5 mm and two to three 1 mm diameter steel grinding beads.
6. Grind in TissueLyser for 5 min. @ 30 Hz.
7. Add 1,000 µL SWB, then spin @ 6,000 *x g* for 5 min. **Important**: work quickly, as some DNA degradation can occur if this step is prolonged. Take note of the supernatant's viscosity and colour. Discard the supernatant and **repeat** the wash if the supernatant was dark and/or viscous.
8. Add 800 µL Extraction buffer, followed by 80 µL 20% SDS.
9. Mix well and place on heat at 65°C for 45 min. to 1 hour, turning every 10-15 min.
10. After lysis, add 80 µL KAc and mix well.
11. Place in -20°C freezer for 10-15 min. **Important**: do not leave in the freezer for longer! The solution needs to cool but must not freeze. Alternatively, the tubes can be placed on ice for a longer period (¿ 30 min.)
12. While waiting, prepare new 1.5 mL Eppendorf tubes and add 600 µL CIA to each.
13. Remove samples from freezer and spin @ 6,000 *x g* for 5 min. Recover 800 µL supernatant into the Eppendorf tubes with CIA.
14. Place in tissuelyser and shake for 1 min. @ 10 Hz (Note: make sure tubes are securely shut! We usually avoid using the outer wells of our tube holders as their lids don't sit as flush on the tube lids at the edges).
15. Spin @ 17,000 *x g* for 8 min.
16. Prepare new 1.5 mL Eppendorf tubes and add 150 µL 5M NaCl and 50 µL Na Acetate to each.
17. Recover *ca.* 700 µL of the aqueous phase into the new tubes.
18. Add 500 µL cold isopropanol and place in -20°C freezer for *ca.* 15 min. **Important**: do not extend this step beyond *ca.* 30 min. as the salts may precipitate.
19. Spin @ 17,000 *x g* for 10 min. Discard liquid and place inverted on two layers of kimwipe. (**Note**: take care not to lose the pellets at this stage!) Allow to dry for 8 min. Turn tubes on their side if the pellet seems loose.
20. Add 1,000 µL 70% EtOH.
21. Flick/shake tubes to loosen pellet.
22. Spin @ 17,000 *x g* for 3 min.
23. Discard ethanol (carefully!) and invert briefly on fresh kimwipe, taking care not to lose the pellet. Seal tube immediately after inversion and place back in centrifuge.
24. Briefly spin down (10 seconds).
25. Pipette out residual ethanol and leave tubes open in rack for 5 min.
26. Check that no droplets remain and elute in 105 µL TE (or as desired – enough for qubit, nanodrop and gel electrophoresis + volume required for sequencing).
